# Subtype-specific circadian clock dysregulation modulates breast cancer biology, invasiveness, and prognosis

**DOI:** 10.1101/2023.05.17.540386

**Authors:** Jan A Hammarlund, Shi-Yang Li, Gang Wu, Jia-wen Lian, Sacha J Howell, Rob Clarke, Antony Adamson, Cátia F. Gonçalves, John B Hogenesch, Qing-Jun Meng, Ron C Anafi

## Abstract

Studies in shift workers and model organisms link circadian disruption to breast cancer. However, molecular rhythms in non-cancerous and cancerous human breast tissues are largely unknown. We reconstructed rhythms informatically, integrating locally collected, time-stamped biopsies with public datasets. For non-cancerous tissue, the inferred order of core-circadian genes matches established physiology. Inflammatory, epithelial-mesenchymal transition (EMT), and estrogen responsiveness pathways show circadian modulation. Among tumors, clock correlation analysis demonstrates subtype-specific changes in circadian organization. Luminal A organoids and informatic ordering of Luminal A samples exhibit continued, albeit disrupted rhythms. However, CYCLOPS magnitude, a measure of global rhythm strength, varied widely among Luminal A samples. Cycling of EMT pathway genes was markedly increased in high-magnitude Luminal A tumors. Patients with high-magnitude tumors had reduced 5-year survival. Correspondingly, 3D Luminal A cultures show reduced invasion following molecular clock disruption. This study links subtype-specific circadian disruption in breast cancer to EMT, metastatic potential, and prognosis.

## Highlights

- Breast cancers exhibit subtype-specific and estrogen-dependent clock disorganization.
- Luminal A tumors show dysregulated rhythmic pathways and varied rhythm strength.
- Higher rhythm strength in Luminal A tumors was correlated with reduced 5-year survival.
- Reducing rhythm strength in Luminal A cells in vitro slows cell invasion.

## Introduction

Worldwide, breast cancer is the most common cancer among women^1–3^. Over the last decades, the introduction of early detection programs, combined with improvements in systemic therapies, have reduced breast cancer mortality^2–4^. Despite these improvements, resistance and subsequent relapse remain major issues^3^. Among women, those with breast cancer lose more disability-adjusted life years than with any other cancer^5^. Adverse effects frequently compromise quality of life^3, 6–8^. There remains a clear need to improve the therapeutic index for breast cancer treatments.

Over the last two decades, research has highlighted the critical roles of cell-intrinsic circadian rhythms in disease (including cancer) and medicine^9–12^. The circadian (∼24-hourly) clock is evolutionarily ancient and highly conserved, permitting cells to anticipate daily environmental changes through temporally coordinated metabolic and gene expression profiles^13–15^. A series of transcription-translational feedback loops form the molecular circadian clock^13–16^. The positive arm of the central loop includes the transcriptional activators Circadian Locomotor Output Cycles Kaput (CLOCK) and Brain and Muscle ARNT-Like 1 (BMAL1)^13–16^. The negative arm, which includes the Cryptochrome (*Cry1/Cry2*) and Period (*Per1/Per2*) genes, later represses translation^13–16^.

The influence of circadian time on cell division, and, by extension, cancer, is particularly strong. In yeast and mammals, evolutionary pressures acted to gate DNA replication and cell division, shielding the dividing cell from solar radiation and toxic metabolites^12, 17, 18^. The cell cycle and circadian clocks share components and signaling molecules and show reciprocal regulation^19–24^. In addition, several oncogenes have been causally linked to circadian clock dysfunctions and may directly hijack clock mechanisms^22, 25^.

Epidemiological and animal studies have proposed that night shift work that disrupts circadian rhythms may increase the risk of developing breast and other cancers^26–28^. This has prompted the WHO to classify night shift work as a probable carcinogen^26–28^. Previously we demonstrated that the extracellular microenvironment modulates functional, cell-intrinsic circadian clocks in mouse mammary gland tissue and human breast epithelial cells^29–32^. Time series transcriptomic analysis revealed hundreds of rhythmic genes in the mouse mammary gland. Rhythmic transcripts included critical molecules implicated in cell cycle regulation, epithelial/progenitor cell function, and hormone responsiveness. Acting, in part, through an immunosuppressive shift in the tumor environment, chronic circadian disruption increases mammary cancer cell dissemination and metastasis in a mouse model of tumorigenesis^33^.

Beyond this basic biology, the molecular circadian clock regulates thousands of genes in a cell and tissue-specific manner. Half of the 100-best-selling-drugs target molecules that oscillate in different mouse tissues^16^. In 1973, Halberg and colleagues initiated a series of studies with the underlying hypothesis that dosage time could influence chemotherapeutic pharmacokinetics, limit toxicity, and improve efficacy^34^. Decades of clinical experience show that time-of-day can influence chemotherapeutic activity^35^. However, the widespread translation of circadian biology in oncology remains slow and serendipitous. Mechanistic knowledge about the unique molecular rhythms in distinct tumors and normal human tissues must be improved. Repeated biopsies or time course sampling from large numbers of human patients is neither safe nor practical. As a result, clinically relevant molecular rhythms still need to be discovered, and opportunities for targeted circadian therapies are unrealized. In addition, while some in vitro and in vivo cancer models demonstrate a complete lack of rhythms, other models (like U2OS cells) show continued rhythms^19^. Indeed, informatic analysis of intact hepatocellular carcinoma has shown disrupted yet persistent transcriptional rhythms^36^.

We recently optimized CYClic Ordering by Periodic Structure (CYCLOPS) to overcome this problem. This machine learning algorithm has allowed us to reconstruct circadian rhythms in samples where sampling time is unknown^36–38^. We adapted CYCLOPS to better account for the non-circadian variation inherent to large-scale data in clinical databases (CYCLOPS 2.0). We then adopted a hybrid study design (Fig 1A). We combined deep sequencing of a small number of time-stamped, paired clinical samples with data from large RNASeq datasets—the Tissue Cancer Genome Atlas^39^ (TCGA) and the Genotype-Tissue Expression^40^ (GTEx) project—where the circadian time of sample collection is unknown. Both the modified CYCLOPS algorithm and experimental validation reveal clear cancer-subtype-dependent changes in molecular clocks and their rhythmic targets. Notably, we uncover a key role of molecular timekeeping in Luminal A tumors linking circadian rhythms to epithelial-mesenchymal transition (EMT), cell invasion, and prognosis. Our studies also provide mechanistic insights into the role of estrogen receptors (ER) in regulating breast cancer clocks.

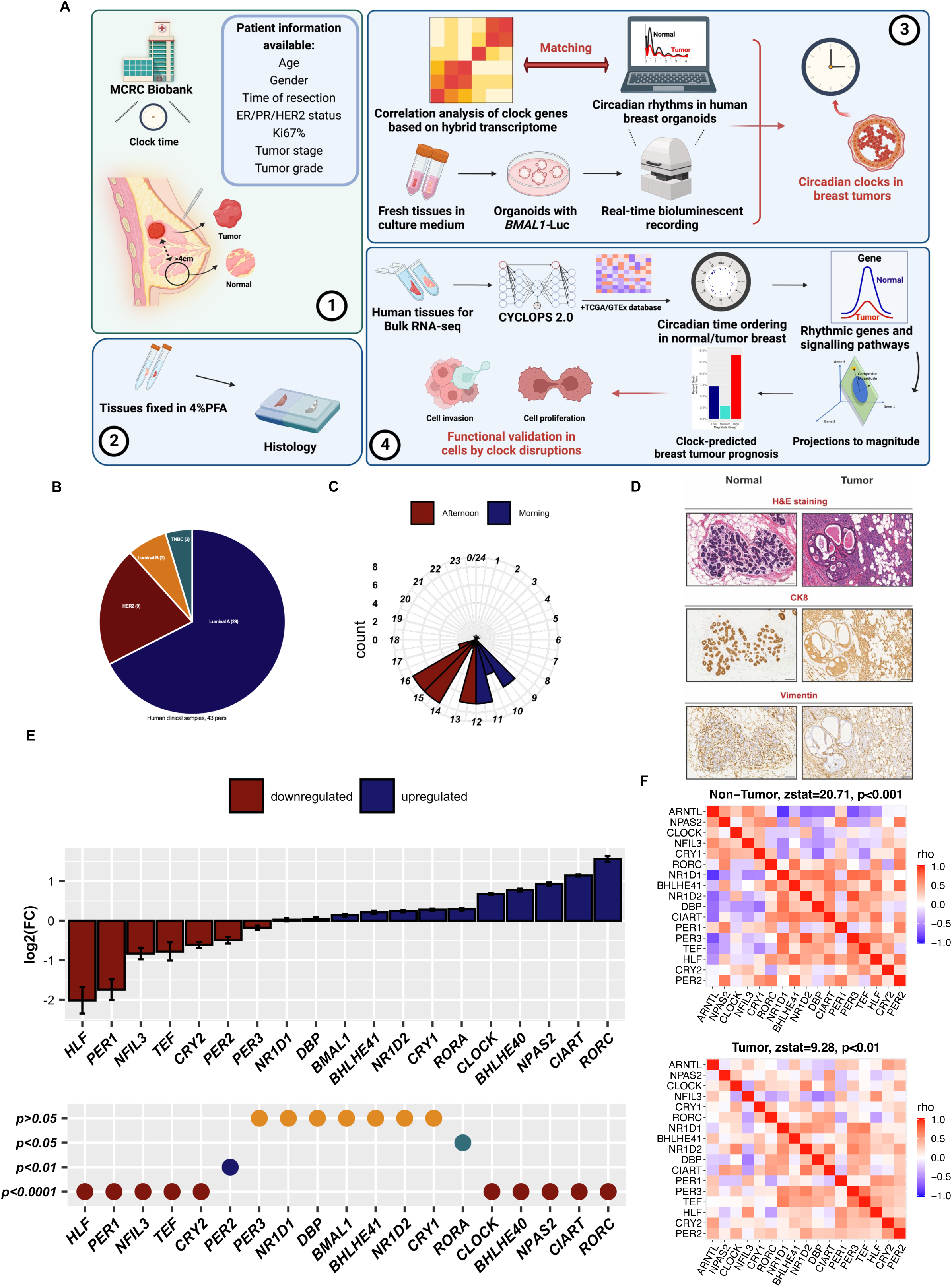
Fig 1.

## Results

### Profound changes in clock gene expression and circadian organization in time-recorded breast cancer biopsies

To assess and improve the accuracy of informatic predictions and to enable direct comparisons of transcriptional changes in circadian genes, we collected 43 pairs of fresh human breast samples (non-cancerous and paired tumors from the same individuals) from patients undergoing mastectomy at the Nightingale Breast Centre, Manchester, UK (Patient demographics on Table. 1). Non-cancerous tissues were collected at least 4 cm away from tumors (Fig. 1A). Tumor samples included Luminal A (N=29), Luminal B (N=3), HER2 (N=2) and Triple-negative breast cancers (TNBC, N=9) (Fig. 1B). Resection times of all samples were recorded (Fig. 1C). To identify non-cancerous and tumor areas we employed Hematoxylin and Eosin-Y staining (H&E staining) and immunohistochemistry using epithelial and stromal markers (Cytokeratin 8 and Vimentin, respectively) (Fig. 1D). Normal breast contains organized acinar and lobular structures, whereas tumor regions lack regular glandular structures. We performed RNA sequencing following RNA isolation. Based on these RNAseq data, the expression of most clock genes is significantly altered in breast cancer tissues. Compared to the paired non-cancerous samples, we observed significant downregulation of PER1, PER2, CRY2, HLF, TEF, and NFIL3 (Fig. 1E; Fig. S1). In contrast, CLOCK, NPAS2, CIART/CHRONO, BHLHE40, RORA, and RORC were significantly upregulated in these breast tumors (Fig. 1E; Fig. S1).

To examine core clock organization in these breast tumors, we performed Spearman’s correlation coefficient analysis^38, 41–43^ using RNAseq data from time-stamped paired breast tumors and non-cancerous samples with a sequencing depth of >20 million reads. As expected, the non-cancerous breast tissues demonstrate core-clock correlation patterns mirroring those seen in mice^16^. The expression levels of clock activators positively correlate with each other across samples. The same is true for the canonical repressors. In contrast, the two groups negatively correlate with each other (Fig. 1F). The strong similarity between the clock correlation patterns seen in the non-cancerous tissue (Fig. 1F, Zstat score of 20.71) and the mouse model suggests a functional clock network. In contrast, a weaker overall correlation in breast tumors (Fig. 1F, Zstat score of 9.28) suggests a weakening of core circadian organization in these samples.

### Transcriptional circadian rhythms are evolutionarily conserved in non-cancerous human breast and mouse mammary tissue

We adopted a hybrid study design^38^ to evaluate circadian time order in human non-cancerous breast tissues. We used informatic tools to integrate RNAseq data from newly collected time-stamped breast samples (N=26, with non-cancerous samples with > 20 million reads) with RNAseq data from female breast samples in public databases. We incorporated data from TCGA^39^ and GTEx^40^ (Table S1). We did not include samples collected in centers where only a small number (n<5) of non-cancerous samples were processed.

Systematic differences between sample collection sites, processing methods, and patient populations complicate the use of aggregate data. These differences may be particularly problematic when different centers have different biases in collection time. We modified the CYCLOPS^36^ neural network to accommodate explicit confounding variables, simultaneously learning confounder adjustments and a common circular structure that explains the variance of the combined data: CYCLOPS 2.0 (Fig. S2A). We benchmarked CYCLOPS 2.0 on actual, semi-synthetic, and fully-synthetic data with different temporal biases (Fig. S2B-D). CYCLOPS 2.0 demonstrates improved accuracy with realistic levels of non-circadian noise. Finally, we allowed the ordering process to use information from a subset of time-stamped samples. We performed 10-fold cross-validation to determine the relative weight given to predicting time in these samples and identify a common circular structure for all samples.

We identified the human orthologues of transcripts that cycle in mouse mammary gland tissue^29^. Combining these with the human orthologues of transcripts that cycled in >75% of mouse tissues^16^, we constructed a circadian “seed gene” list appropriate for ordering human breast tissue. Ordering the combined dataset using these seed genes and including temporal information from the 26 time-stamped human samples, the CYCLOPS smoothness and ordering metrics for the entire dataset meet previously established standards (Statsmooth=0.75, Staterror=0.015). The CYCLOPS-predicted sample phases show a significant correlation with the known sample collection times of the 26 subjects (Corrcirc=0.7, p<0.005) (Fig. 2A). As expected, the clinical biopsies available in TCGA show a temporal bias in inferred sample collection phase (Fig. 2B). In contrast, the distribution of inferred sample phases assigned to the GTEx samples (autopsy collection) was more uniform (Fig. 2B). After ordering, we used modified Cosinor regression^36, 44^ to identify cycling transcripts and estimate their amplitude and acrophase (time of peak expression) (Fig. 2C, D). The expanded Cosinor model explicitly accounted for differences in expression due to sequencing sites or source databases. At a BHq threshold of 0.05, we identified ∼2,000 genes as rhythmic. When we imposed a relative cycling amplitude (amplitude/MESOR (Midline Estimating Statistic of Rhythm)) greater than 1/3— as a measure of likely biological significance—we reduced the number of identified cycling transcripts to ∼650 (File. S2). As observed in other tissues, there are clear circadian “rush hour periods” where many rhythmic transcripts peaked^16, 37, 45^. With the notable exception of RORC, the relative acrophases of core-clock transcripts reconstructed from non-cancerous human breast tissue are in good accord with the well-established ordering of these transcripts in other mouse and human tissues (Fig. 2E).

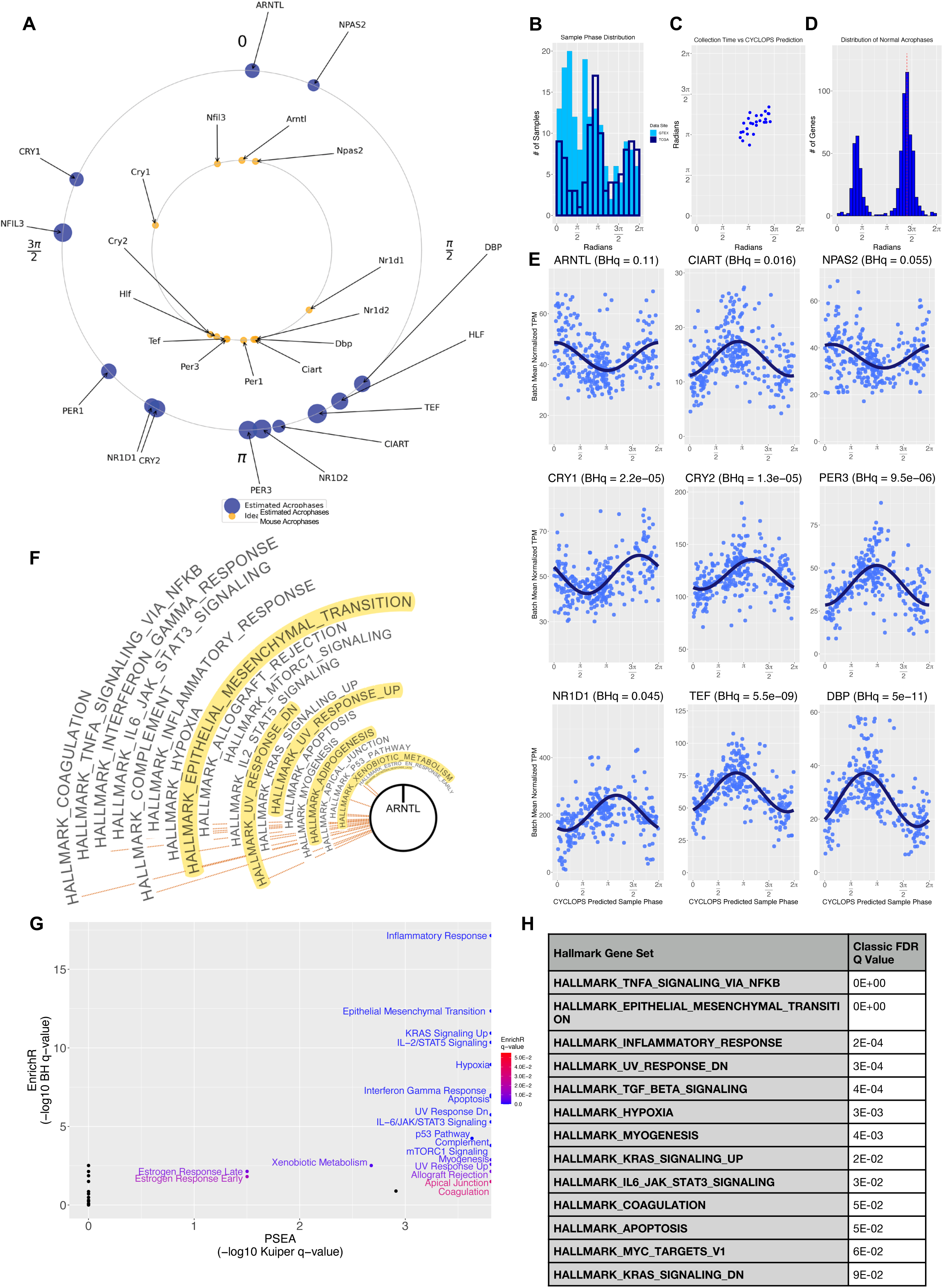
Fig 2.

To put these cycling results in a broader biological context, we used phase set enrichment (PSEA)^46^ to identify annotated gene sets and biological pathways where the constituent cycling transcripts exhibited circadian concentration and were not uniformly distributed across the circadian day. Pathways related to adipogenesis, EMT, and estrogen responsiveness, show circadian orchestration, similar to reports from mouse mammary gland (Fig. 2F). Using both gene set enrichment^47^ and over-representation approaches^48–, 50^, we also labeled pathways that were enriched for cycling genes. In addition to the abovementioned pathways, various immune and cell cycle pathways show marked circadian orchestration (Fig. 2G, H).

### ER activity correlates with circadian organization and function in breast cancer subtypes

Our clock correlation analysis on locally collected cancer samples combined data from biologically distinct breast tumor types. We applied clock gene correlation analysis^38, 41–43^ to TCGA breast tumor data to evaluate cancer-subtype-dependent changes in clock organization—the expression of the PAM50 panel genes defined cancer subtypes^51, 52^. Consistent with the non-cancerous time-stamped samples, the non-cancerous breast tissues from the database show an intact core circadian organization that closely mirrors the established consensus with a Zstat score of 20.86. The Luminal A samples demonstrate weaker but still considerable evidence of intact circadian organization with a Zstat score of 11.04. On the other hand, Luminal B and Triple Negative breast cancers exhibit disrupted correlation patterns with Zstat scores of 6.93 and 4.98, respectively (Fig. 3A). The relatively small number of HER2 samples in the TCGA database prevented evaluation of clock organization in this tumor type.

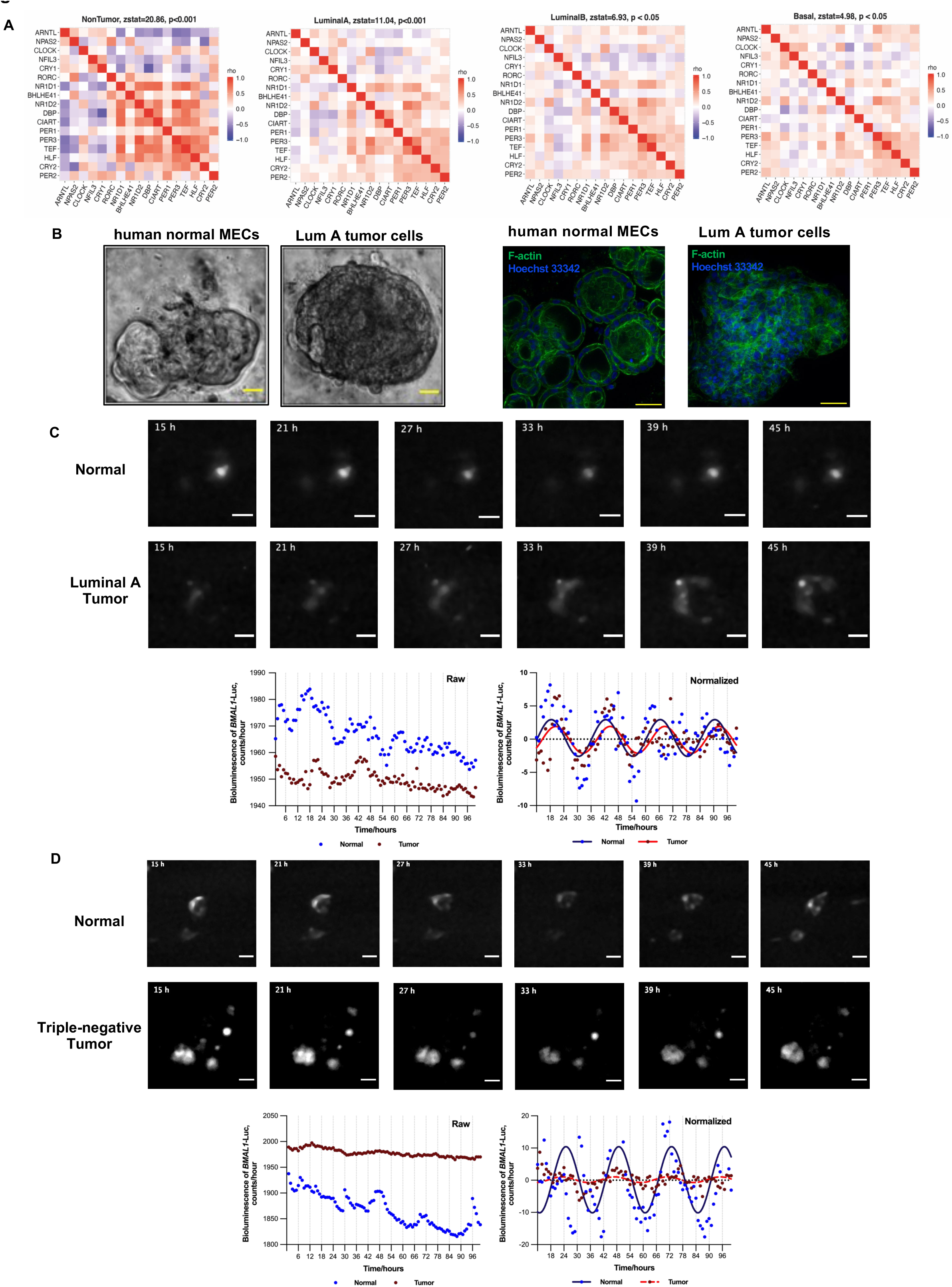
Fig 3.

Given these findings, we hypothesized that circadian function in breast tumor tissues, like core clock organization, varies among cancer subtypes - with Luminal A samples showing reasonably good clocks. We assessed molecular circadian rhythms in breast tumors and paired non-cancerous tissues from the same individuals to confirm these predictions. We derived organoids from primary mammary epithelial cells. This model more closely mimics in vivo physiological functions of mammary epithelia^31, 32, 53^. The organoid cultures from normal breast tissues showed typical acinar structures. In contrast, breast tumor organoids showed disrupted cell polarity (Fig. 3B). After lentiviral transduction of a BMAL1-Luc circadian reporter, we imaged bioluminescence signals using an LV200 imaging system (Olympus). Non-cancerous human mammary organoids showed robust circadian rhythms. Patient-derived Luminal A tumor organoids showed persistent but weakened rhythms (Fig. 3C, Video. S1, N=4). However, we did not observe sustained circadian rhythms in BMAL1-Luc activity in TNBC tumor organoids (Fig. 3D, Video. S2, N=3), Luminal B organoids, or HER2 organoids (data not shown). Using either a BMAL1-Luc reporter or time course western blot studies of clock factors, we also observed corresponding clock changes in established breast cancer cell lines representing various tumor subtypes (Fig. S3A-C). The MCF-7 cell line (representative of ER+ Luminal A) continues to exhibit circadian rhythms, while the MDA-MB-231 (TNBC) and SKBR3 (HER2+) cell lines do not (Fig. S3B, C). In unsynchronized cells, we observed altered average expression levels of core clock genes among the three cell lines. (Fig. S3D).

As ER status is one of the key factors in differentiating these tumor subtypes, our informatic and experimental results suggest a possible link between ER responsiveness and clock functions in breast cancer cells. Indeed, further stratification of breast tumor samples based on ER status indicates a strong correlation between ER expression and clock functionality (Fig. S4A). To more directly determine whether ER signaling regulates circadian rhythms in breast cancer cells, ERα was knocked out of BMAL1-Luc MCF-7 cells using CRISPR-Cas9. Following co-transfection of sgRNA and Cas9 protein, single-cell colonies were isolated. We confirmed successful knockout by DNA sequencing, supported by the absence of ERα mRNA and protein (Fig. S4B, C). ERα-KO disrupted the expression of clock factors in MCF-7 cells compared to the control (Fig. S4 D, E). In contrast to the robust 24-hour rhythms in control MCF-7 cells, there was a complete loss of circadian BMAL1-Luc rhythms in all four clones of MCF-7 cells with ERα-KO (Fig. S5A). In addition, an ERα selective agonist PPT (Propyl Pyrazole Triol) synchronized circadian clocks in MCF-7 cells in a dose-dependent manner, further supporting a regulatory role of ER signaling in MCF-7 cell circadian function (Fig. S5B).

### CYCLOPS 2.0 analysis revealed global changes in rhythmic gene expression patterns and pathways in Luminal A samples

Guided by the experimental and informatic evidence for persistent rhythms in Luminal A tumors and the relative abundance of Luminal A samples in the TCGA database, we next used CYCLOPS 2.0 to order Luminal A tumors (Table S2). There is likely significant non-circadian heterogeneity among Luminal A samples. We projected the Luminal A data onto the eigengene space computed from the non-cancerous samples^36^ to emphasize the variation resulting from circadian time. Using the CYCLOPS 2.0 model, we included data from both non-cancerous and Luminal A samples in the ordering, now listing tumor status as a covariate. After ordering and applying cosinor regression to the Luminal A samples, seven core clock genes, including DBP, NR1D1, NR1D2, TEF, PER3, NFIL3, and CRY1, meet the initial criteria for cycling (Fig. 4A, B). At a BHq threshold of 0.05, we identified ∼1,100 genes as rhythmic. When we imposed a relative cycling amplitude greater than 1/3, we reduced the number of identified cycling transcripts to ∼675 (File. S2). Of course, differences in sample size and non-circadian variability may have contributed to these changes. Thus, in addition to simply identifying genes that meet our statistical cutoffs in Luminal A, we used nested regression models as we^36^ and others^54, 55^ have done previously, to directly test for changes in cycling between Luminal A and non-cancerous samples (File. S3). This nested modeling approach tests the importance of tumor-dependent cycling parameters while accounting for tumor-dependent differences in mean expression level. We observe changes in core clock gene and clock output gene rhythms in Luminal A samples (Fig. 4A, B). For example, while TEF shows decreased amplitude in the Luminal A samples, its partner and structural/functional paralogue^56^, DBP, shows increased amplitude. As in the non-cancerous samples, Luminal A samples show “rush hours” of rhythmic transcription (Fig. 4C). However, here, the proportion of samples assigned to the window that precedes the ARNTL (BMAL1) acrophase (inferred early evening) is much higher. Using our nested regression models to compare the fit amplitude for transcripts that cycled in either Luminal A or non-cancerous samples, we find that more transcripts lose as opposed to gain amplitude in Luminal A samples (Fig. 4D). Of note, TCGA includes a limited number of matched Luminal A tumors and non-cancerous samples from the same patient. The sample phases assigned to the tumors and their non-cancerous matches are poorly correlated (Fig. 4E).

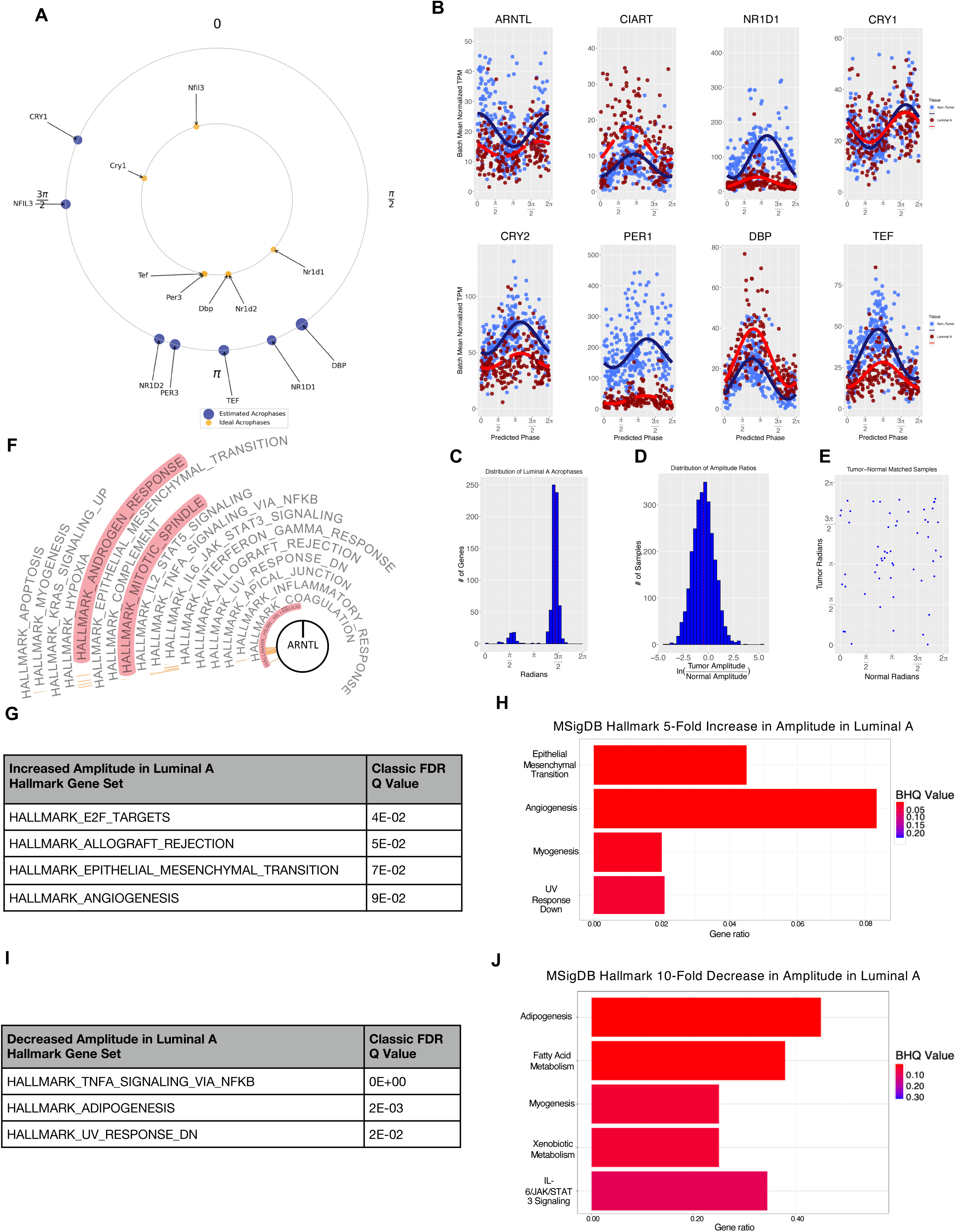
Fig 4.

At a gene set level, many pathways demonstrate continued circadian orchestration in the Luminal A samples (Fig. 4F). Phase set enrichment analysis reveals cycling in EMT, androgen responsiveness, and numerous immune and inflammatory pathways. We used two complementary approaches to focus on pathway-level differences in Luminal A and non-cancerous output rhythms. To identify pathways with enhanced rhythmicity in Luminal A, we first ranked the full complement of genes that cycled in either Luminal A or non-cancerous tissue by the log fold change in amplitude. We then used GSEA to identify gene sets enriched for more marked amplitude increases in this ranked list. Alternatively, we used nested regression to identify transcripts that showed statistically significant differential cycling (BHq<0.05) between Luminal A and non-cancerous samples. Focusing on genes with a more than five-fold gain in amplitude in Luminal A samples, we used EnrichR to identify pathways overrepresented in this discrete set. Both methods yield similar results (Fig. 4G, H). EMT and angiogenesis pathways—critical to cell invasion and growth support—show increased cycling in Luminal A samples. We find that adipogenesis appears to have reduced cycling using the same two analyses (Fig. 4I, J). Pathways related to fatty acid metabolism and NFƙB signaling (among others) also show reduced cycling in Luminal A tumors using one or the other analysis (Fig. 4I, J).

### CYCLOPS magnitude as a measure of global circadian rhythm strength

While we understand the amplitude of a single rhythmic waveform, meaningful, global measures of transcriptional rhythm strength still need to be well established. For example, it is generally unknown if high amplitude circadian expression in some genes predicts high amplitude circadian expression of others. CYCLOPS operates on eigengenes—global descriptors of expression that summarize the behavior of many cycling genes. CYCLOPS projects these data onto a plane where a circular structure is apparent. We use the angular position of any sample on this circle to infer its internal molecular phase. We also calculate the radial distance of each sample from the circle’s center. Geometrically, we interpret CYCLOPS magnitude (CMag) (Fig. 5A) as a weighted sum of the amplitudes of the individually cycling seed genes. This concept resembles the PCA plots of cycling gene expression in Brooks et al.^57^. The distribution of CYCLOPS magnitudes obtained from the Luminal A samples is broad with a long tail (Fig. 5B). Dividing samples into equal thirds based on CMag, we find that across all transcripts cycling in Luminal A samples, the amplitude of cycling is generally greater in high magnitude samples as compared to low magnitude samples (Fig. 5C, D). Unlike Luminal A samples as a whole (Fig. 4E), the circadian molecular phases assigned to high CMag Luminal A samples generally match the phases assigned to their non-cancerous pair (Fig. 5E). This suggests that higher CMag in Luminal A samples is indicative of a more robust clock.

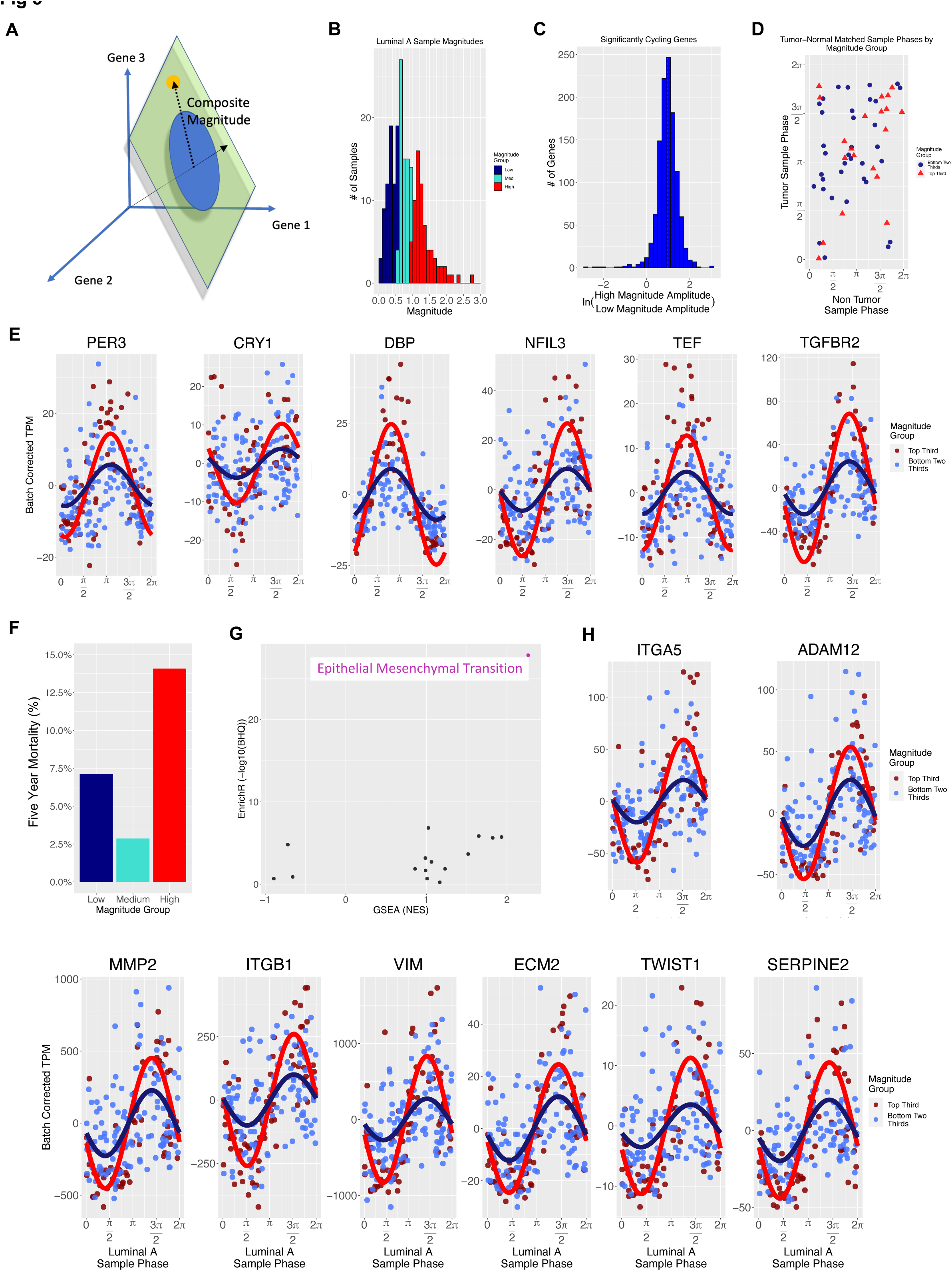
Fig 5.

### Circadian rhythm strength predicts prognosis and modulates metastatic potential

We investigated if rhythm strength influenced tumor biology and prognosis. Patients with Luminal A tumors, as assessed by the PAM50 panel^52^, were stratified into three equally sized groups based on the CMag of their tumors (low, medium, and high CMag). Using TCGA outcome data, we evaluated these patients’ 5-year survival. Retrospectively, the risk of death increases among patients with high-magnitude Luminal A tumors (Fig. 5F). This difference is statistically significant (p=0.047, ANOVA). The increased risk in the high-magnitude tumor group remains statistically significant when we tested its influence in a generalized (logistic regression) model that also included patient age and the presence of known metastases at diagnosis (p<0.05). A high-magnitude tumor increased the relative risk by ∼1.5x, over and above the risk established by the other covariates. Indeed, the predictive value of tumor magnitude remains significant in a model that includes MKI67 transcript expression (p<0.05). Of some note, while the same trend appears examining a broader outcome of death OR new cancer event (Fig. S6), this trend is not statistically significant.

CYCLOPS magnitude is broadly associated with the cycling amplitude of many genes. Given its prognostic importance, we next identified rhythmic pathways that showed the most marked differences in high-magnitude samples. For each transcript that showed statistically significant cycling in Luminal A samples, we compared the amplitude estimated from the high-magnitude samples (top third) to the amplitude estimated from lower-magnitude samples (bottom two-thirds). Again, we leveraged both enrichment and overrepresentation approaches to analyze these results at the pathway level. When we compare high-and lower-magnitude samples, our analyses show that EMT-related genes exhibit the most pronounced changes in cycling (Fig. 5G, H). Given the well-established role of EMT in tumor biology and, in particular, metastatic potential, we hypothesized that high amplitude rhythms might modulate Luminal A tumor cell behavior and the potential for invasion.

Most core circadian clock genes have paralogues that can functionally compensate for molecular knockdown^58^. Only BMAL1 is essential for circadian locomotor function^59^. To establish the role of circadian clocks in regulating breast cancer cell behavior, we used lentiviral shRNA for BMAL1 to disrupt cellular clock functions in rhythmic MCF-7 cells. As expected, the lack of BMAL1 abolished circadian reporter rhythms in MCF-7 cells (Fig. 6A). We used hanging drop and cell invasion assays to evaluate the invasion of MCF-7 cells through a 3D collagen I matrix microenvironment. We assessed invasiveness by measuring the distance from the center of the spheroid (initial droplet) to the edge of the furthest cell. Circadian disruption through BMAL1 deficiency inhibited the rate of cell invasion in both MCF-7 cells (p< 0.001, Fig. 6B, C) and primary Luminal A breast tumor cells (Fig. S7A). When we disrupted the molecular clock with KL001, which stabilizes both CRY1/CRY2, we observed a similar suppression of cell invasiveness (Fig. 6D-F). Next, MCF-7 cell proliferation was assessed by expression of Ki-67 and real-time quantitative live cell imaging using IncuCyte. BMAL1 knockdown increased levels of Ki-67 (Fig. S7B) and cell proliferation (p< 0.0001) (Fig. S7C, Video. S3) in MCF-7 cells. As such, the loss of molecular clock rhythm in MCF-7 cells compromises breast cancer cell invasion into the 3D matrix, despite increasing cell proliferation.

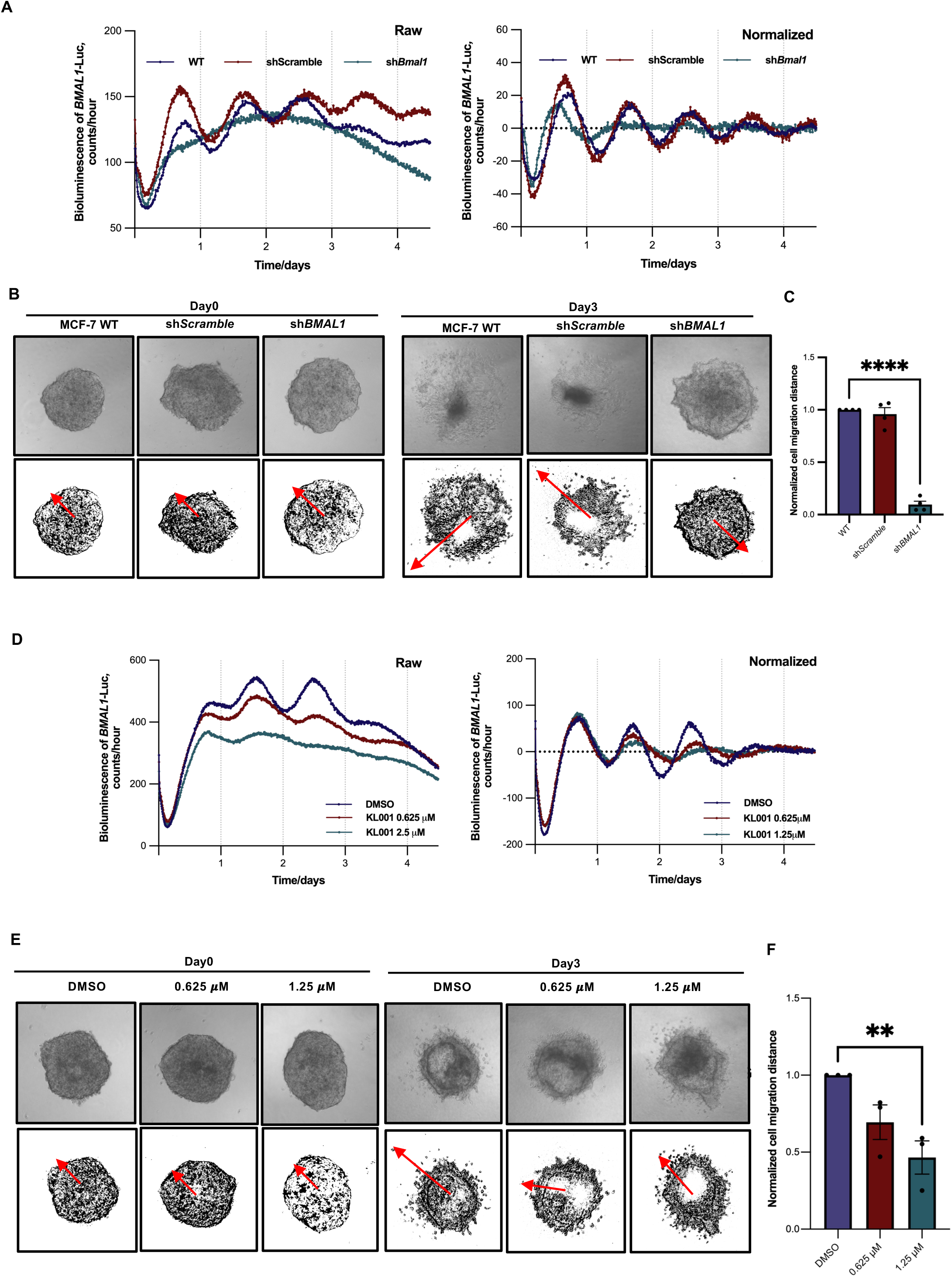
Fig 6.

## Discussion

This work used informatic ordering (CYCLOPS 2.0) to integrate newly collected, time-stamped biopsies with public data. We reconstructed temporal rhythms in non-cancerous breast tissue and Luminal A breast tumors. In non-cancerous tissue, our approach reveals the cycling of inflammatory, EMT, and estrogen response pathway genes. Experiments with Luminal A organoids show continued, albeit dampened rhythms. Disrupted rhythms are also evident in our informatic circadian reconstruction of Luminal A tumors. Strikingly, retrospective analysis shows that Luminal A cancer patients with high rhythm strength tumors had increased 5-year mortality. EMT pathway genes show the most marked increase in cycling when comparing tumors with higher and lower rhythm strength. Given the importance of EMT in cell invasion, we hypothesized that tumors with high rhythm strength might show increased metastatic potential. Accordingly, 3D culture experiments using established Luminal A cancer cell lines and primary Luminal A cells show reduced invasion following molecular clock disruption. As such, our study links subtype-specific circadian disruption in breast cancer to EMT, metastatic potential, and prognosis.

A bi-directional web of transcription factors and direct protein-protein interactions couples the cell-intrinsic circadian clocks and the cell cycle. Core circadian clock genes include the likely tumor suppressors PER1 and PER2^60–62^. More recently, researchers have shown that the master clock activators BMAL1 and CLOCK have anti-apoptotic roles, promoting liver cell proliferation through the cell cycle regulator Wee-140^63^. On the other hand, oncogenes such as c-MYC or KRAS interfere with circadian pacemaking^24, 25^.

The circadian-cancer connection may be vital in breast cancer. Several epidemiologic studies have now linked night shift work with breast cancer risk^26–28^. In mouse models bearing primary mammary tumors or breast cancer xenografts, the efficacies of Doxorubicin and Celecoxib are time-of-day dependent^64, 65^. However, the difficulty of obtaining time-course clinical samples across multiple circadian cycles in a large, clinically informative cohort hinders our understanding of circadian biology and its translation in human breast cancer.

Several supervised learning algorithms (e.g., BodyTime, TimeSignature, TimeTable, and ZeitZeiger) predict internal clock time from time-unknown human samples^66–69^. These approaches require “training data” that spans the tissues and conditions covered in later applications. BodyTime, for example, uses a small set of transcriptomic biomarkers from blood monocytes to predict the melatonin phase^67^. These approaches show promise when applied to the specific tissue for which they were trained. However, they are not designed for application to new tissues or disease states (such as solid tumors) without specific training data.

In contrast, CYCLOPS^36^ uses global descriptors of expression structure and unsupervised machine learning that identifies more general signatures of rhythmic processes. CYCLOPS does not require exact knowledge of the particular cycling genes in a tissue or the specific temporal relationship between those rhythmic genes. Instead, CYCLOPS requires a list of “seed genes” likely to cycle in a given tissue. It assumes a fixed relative phase relationship between the cycling seed genes across subjects (e.g., BMAL1 precedes NR1D1 by a relatively fixed amount in the oscillations in each sample). More recently, other unsupervised and semi-supervised approaches have emerged. Most notably, Talamanca et al.^45^ aimed to increase the power of this approach, focusing on GTEx data, where many sample tissues were taken from the same individuals. Their approach assumes that tissues obtained from the same individual at the same time are at the same molecular phase.

In our work, however, the limitations of having a single tissue type are compounded by the need to aggregate data from several sources. We specifically tailored our modifications to CYCLOPS for these issues. In this context, non-circadian covariates and batch effects in processing can likely overwhelm circadian variation. As we demonstrate in our benchmarking, batch-correcting approaches like COMBAT that attempt to “normalize away” these differences are unlikely to overcome this obstacle. If different centers have different biases in collection time, that approach may remove much of the circadian signal. This challenge is particularly relevant when combining clinical biopsy and autopsy-based collections. CYCLOPS 2.0 explicitly accommodates these issues, finding batch and covariate adjustments to utilize a common underlying periodic structure for all datasets.

While our CYCLOPS 2.0 ordering of non-cancerous breast tissue is consistent with well-established circadian physiology (i.e., the relative phase relationships between core clock genes) and meets various informatic quality checks, it is always reasonable to question informatic results. We cannot dismiss the influence of non-circadian variability on the ordering process. Our hybrid experimental design lends considerable reassurance. This design required only a small number of newly collected time-stamped samples to guide our efforts and demonstrate that the informatic ordering reflects natural temporal variation. This informatics-guided approach reflects a practical compromise, allowing us to infer molecular rhythms without exhaustive time course sampling more confidently. It should also be noted that CYCLOPS, like other ordering methods, infers time as a function of gene expression. Therefore, using time as an independent variable for cycling analysis is somewhat fraught. We have attempted to address this, as we did previously^36^, by implementing a more stringent modified cosinor regression and imposing strict numerical thresholds.

Clock gene correlation analysis from breast cancer samples showed subtype-dependent changes in the core-clock organization, supported by our in vitro data showing subtype-dependent clock functionality in tumor organoids. Both approaches revealed a critical role in estrogen responsiveness in regulating breast cancer clocks. Previous work using microarray data to study the correlation between clock genes in node-negative breast cancer patients supports this concept of breast cancer subtype-dependent clock changes. Pairwise correlations between functionally related clock genes (e.g., PER2-PER3 and CRY2-PER3) were more robust in ER+/HER2- and weaker in ER-/HER2+ tumors^70^.

Among breast cancer subtypes, Luminal A tumors had the most robust evidence for persistent rhythms, prompting our interest in ordering these tissues along circadian time. We find marked changes in the informatically reconstructed cycling in Luminal A tumors; many genes and pathways, including chemotherapeutic targets, gained or lost rhythmicity. Using CYCLOPS magnitude as a measure of the global rhythm strength in each sample, we identify marked variations in the rhythm strength of Luminal A tumors. The global magnitude of rhythmic oscillation in Luminal A tumors negatively correlates with five-year mortality and positively correlates with cycling in EMT pathway genes. Our in vitro experimental evidence casually links molecular clock disturbance with cancer cell invasion in a 3D model, thus supporting our informatic result. The work of De et al., who observed MCF-7 cells and noted circadian rhythms in EMT-associated changes in cell morphology^71^, also buttresses our results.

Our results also agree with a previous analysis of TCGA data comparing paired tumor and non-tumor samples in 14 cancer types. Specifically, in breast cancer, they report a similar downregulation of PER1, PER2, CRY2, and HLF, while CLOCK, ARNTL(BMAL1), and BHLHE40 levels remained relatively unchanged^72^. However, by including a temporal ordering component in the analysis of Luminal A tumors, we observe changes in rhythmic patterns. For example, we observe that ARNTL(BMAL1) loses rhythmicity in addition to a change in basal expression. Similarly, while HLF cycling shows increased amplitude, its functional paralogue^73^ DBP shows decreased cycling amplitude. This data will allow cancer researchers to identify chemotherapeutic targets with temporal properties which differ between cancer and non-cancerous tissue, opening a potential chronotherapeutic opportunity. For example, we note cycling in BRAF and other kinase pathway target genes (File. S4-S8). We hypothesize that highly rhythmic Luminal A tumors, which have the worst prognosis, likely due to increased invasiveness despite the reduced proliferative potential, will also be most responsive to time-aware therapies.

These new insights bring us one step closer to personalized circadian medicine. Our results also underscore vital outstanding questions. Comparing rhythms in non-tumor and luminal A samples, the EMT pathway genes stood out for demonstrating increased cycling in tumor samples. Comparing cycling between Luminal A tumors with high/low global rhythm strength again highlights the EMT pathway. Our repeated identification of EMT as a rhythmically coordinated pathway in Luminal A tumors is particularly intriguing, given recent reports that the metastatic spread of breast cancer accelerates during sleep^74^. Although our experimental results show that Luminal A tumors have cell-autonomous rhythms, our informatic study design cannot distinguish clinically relevant direct clock outputs from rhythms imparted from cycling hormones or other physiological signals. Tumors with high rhythm magnitudes may be more responsive to host signals. Using mouse models, Hill and colleagues previously identified nocturnal light exposure and the corresponding change in host melatonin rhythms as influencing EMT^75^. While the association between tumor rhythm strength, EMT cycling, and patient prognosis is important regardless of mechanism, distinguishing these possibilities is likely essential for targeted therapy.

Further investigations are needed to evaluate the relative contributions of host and tumor rhythms in modulating human disease. Our experimental results showing a change in tumor invasiveness following clock disruption suggest that tumor-autonomous rhythms causally influence metastatic potential. However, it is also possible that circadian fitness or responsiveness is a marker of other features that contribute to an aggressive phenotype. In addition, we cannot be sure these disrupted tumor rhythms retain a ∼24-hour period in vivo. As CYCLOPS does not explicitly measure time but rather an ordering relative to an internal molecular cycle, it is possible that in vivo tumor-autonomous rhythms have a longer or shorter period.

Of particular note, our result suggesting tumor rhythm strength is a potential prognostic marker requires significant follow-up and prospective verification in an independent cohort before any clinical application should be considered. Currently, CYCLOPS uses the complete list of ∼70 “seed genes” to compute the magnitude score. A smaller cohort of transcripts could likely suffice for this purpose. While circadian magnitude offers prognostic value beyond PAM50 tumor type and MKI67 levels, we do not have immunostained Ki-67 levels in these TCGA samples. While MKI67 transcript and Ki-67 protein levels correlate, they are clearly different. However, we predict that the mechanistic insights our results suggest and the awareness that high-magnitude tumors are better candidates for circadian medicine approaches may prove most useful.

Our results also emphasize the importance of subtype and patient-specific analysis of tumor rhythms. The interactions between cancer biology and circadian rhythms are multifaceted and tumor dependent. The biological differences between tumor subtypes extend far beyond the clock. Indeed, our screening analysis, like previously noted informatic studies, suggests that more aggressive HER2 and triple-negative tumors have weaker or absent rhythms. Nevertheless, we find that within Luminal A tumors, increasing rhythm strength appears to predict increased invasiveness. We believe it is ill-advised to make blanket statements based on a single tumor type or to compare rhythms in tumor types with vastly different biological features. Our results cannot be applied directly to other tumor types. Future studies could test the intriguing hypothesis that specific cancer cells “hijack” the clock mechanism to temporally organize metabolic programs, evade immune surveillance, suppress apoptosis, or facilitate intravasation and metastasis.

Some cancers may specifically disrupt the circadian check on cell division. Other cancer cells may have evolved to break loose from circadian control altogether to suit their needs best^63, 73^. For Luminal A tumors, like most organisms, tumor rhythms may impart increased biological fitness – to the detriment of the patient. Taken as a whole, the biological insights from this study may help lay the groundwork for improved breast cancer prevention (e.g., lifestyle changes), new prognostic biomarkers, and more effective personalized breast cancer treatments.

## Supporting information

Figure Titles and Captions

## Acknowledgments

J.A.H., G.W., J.B.H. and R.C.A. were supported by the National Cancer Institute (5R01CA227485). S-Y.L, S.J.H., R.C. and Q-J.M. were supported by the Breast Cancer Now grant (2022FebPR1518). S-Y.L. and Q-J.M. were also supported by the Helen Muir Fund from the Wellcome Centre for Cell-Matrix Research (088785/Z/09). R.C.A. received additional support from 5R01AG068577. J-W.L. was supported by a Medical Research Council PhD Studentship. We would like to thank the Manchester Cancer Research Centre Biobank for access to clinical breast samples and pathological characterizations. We thank Prof. Charles Streuli for his invaluable inputs and advice with the design of this project. We thank Dr. Dharshika Pathiranage, Miss Xiang-jun Zhao for their kind assistance with various experiments. We also thank the Histology and Bioimaging Core Facilities and the Genome Editing Unit (University of Manchester) for their technical support and advice.

## Author contributions

R.C.A., Q-J.M. and J.B.H. conceived the study. Q-J.M., R.C.A., J.B.H., S.J.H. and R.C. secured funding. S-Y.L., J.A.H., G.W, J-W.L. and C.F.G performed experiments. R.C.A., Q-J.M., G.W., S-Y.L. and J.A.H. wrote the manuscript. All authors edited the manuscript. S.J.H., R.C.A. Q-J.M, and A.A. supervised the project and provided expertise, reagents, and feedback.

## Declaration of interests

J.B.H. is on the scientific advisory board for Synchronicity Pharma. The remaining authors declare no competing interests.

**Tables. 1.** Patient demographics for 43 pairs of Manchester human breast samples

| Patient ID | Gender | Age | Time of resection | Type | Grade | ER<sup>a</sup> | PR<sup>b</sup> | HER2<sup>c</sup> | Ki-67 |
| --- | --- | --- | --- | --- | --- | --- | --- | --- | --- |
| W005787 | Female | 72 | 17:10 | IDC | 3 | 0 | 0 | 3 | 40% |
| W005805 | Female | 74 | 16:35 | IDC | 2 | 8 | 8 | 1 | 5% |
| W005821 | Female | 62 | 15:06 | IDC | 3 | 0 | 0 | 0 | 70% |
| W005830 | Female | 38 | 16:05 | IDC | 3 | 8 | 5 | 1 | 38% |
| W005831 | Female | 78 | 12:55 | IMC | 3 | 0 | 0 | 0 | 38% |
| W005859 | Female | 44 | 15:20 | IDC | 2 | 8 | 8 | 1 | 14% |
| W005863 | Female | 51 | 16:40 | IMC | 2 | 8 | 5 | 1 | 36% |
| W005925 | Female | 65 | 15:50 | IDC | 3 | 8 | 0 | 3 | - |
| W006024 | Female | 93 | 12:45 | IDC | 3 | 2 | 0 | 0 | 62% |
| W006029 | Female | 54 | 13:10 | ILC | 2 | 8 | 7 | 1 | 25% |
| W006034 | Female | 76 | 13:50 | IDC | 2 | 8 | 8 | 1 | 21% |
| W006044 | Female | 60 | 12:30 | IDC | 2 | 8 | 8 | 1 | 28% |
| W006045 | Female | 63 | 11:55 | IDC | 1 | 0 | 0 | 1 | 82% |
| W006053 | Female | 47 | 16:00 | IDC | 3 | 8 | 8 | 0 | 50% |
| W006055 | Female | 75 | 13:50 | IDC | 3 | 2 | 0 | 1 | 63% |
| W006071 | Female | 45 | 13:00 | IDC | 2 | 7 | 7 | 2 | 27% |
| W006074 | Female | 75 | 16:10 | IDC + DCIS | 2 | 8 | 7 | 1 | 17% |
| W006077 | Female | 40 | 10:40 | IDC | 3 | 0 | 0 | 0 | - |
| W006078 | Female | 70 | 16:45 | IDC | 2 | 8 | 0 | 2 | 25% |
| W006140 | Female | 77 | 10:25 | IAC | 2 | 0 | 0 | 1 | 12% |
| W006147 | Female | 73 | 15:40 | IDC | 3 | 0 | 0 | 1 | 30% |
| W006155 | Female | 68 | 11:18 | IDC | 3 | 8 | 3 | 1 | 21% |
| W006161 | Female | 60 | 15:55 | ILC | 2 | 8 | 6 | 2 | 10% |
| W006167 | Female | 85 | 10:54 | IDC | 2 | 8 | 7 | 1 | 30% |
| W006193 | Female | 54 | 13:50 | IDC + DCIS | 2 | 8 | 7 | 2 | 36% |
| W006195 | Female | 33 | 11:45 | IDC | 1 | 8 | 7 | 1 | 20% |
| W006212 | Female | 42 | 12:55 | IDC | 2 | 8 | 0 | 3 | - |
| W006217 | Female | 48 | 10:45 | IDC | 2 | 7 | 8 | 2 | 20% |
| W006218 | Female | 76 | 13:15 | IDC | 2 | 8 | 8 | 1 | 12% |
| W006219 | Female | 71 | 12:20 | ILC | 2 | 8 | 4 | 1 | 5% |
| W006226 | Female | 43 | 15:00 | IDC | 3 | 0 | 0 | 3 | - |
| W006238 | Female | 56 | 10:05 | IDC | 3 | 8 | 4 | 2 | 29% |
| W006241 | Female | 80 | 16:45 | IDC | 3 | 8 | 7 | 2 | 35% |
| W006267 | Female | 71 | 17:03 | IPLC | 3 | 7 | 5 | 1 | 30% |
| W006268 | Female | 41 | 12:00 | ILC | 2 | 7 | 7 | 2 | 19% |
| W006312 | Female | 63 | 15:58 | IDC | 2 | 8 | 7 | 1 | 13% |
| W006314 | Female | 54 | 12:15 | IDC | 2 | 8 | 8 | 1 | 19% |
| W006335 | Female | 45 | 11:40 | IDC | 3 | 8 | 7 | 0 | 10% |
| W006506 | Female | 69 | 15:55 | ILC | 2 | 6 | 0 | 2 | 20% |
| W006510 | Female | 49 | 16:20 | IDC | 3 | 8 | 3 | 0 | 33% |
| W006511 | Female | 53 | 10:15 | IDC | 1 | 8 | 6 | 1 | 25% |
| W006542 | Female | 39 | 13:50 | ILC + DCIS | 2 | 8 | 8 | 1 | 10% |
| W006553 | Female | 72 | 14:40 | IDC | 3 | 0 | 0 | 1 | 18% |

a, estrogen receptor; b, progesterone receptor; c, human epidermal growth factor receptor 2

## Key resources table

| REAGENT or RESOURCE | SOURCE | IDENTIFIER |
| --- | --- | --- |
| **Antibodies** |  |  |
| Alexa Fluor™ 488 Phalloidin | Thermo Scientific | Cat# R37110 |
| Anti-Rabbit BMAL1 | Homemade (Weaver Lab) | N/A |
| Anti-Rabbit Cytokeratin 8 | Abcam | Cat# ab53280<br>RRID: AB_860091 |
| Anti-Rabbit CLOCK | Proteintech | Cat# 18094-1-AP<br>RRID: AB_2878497 |
| Anti-Rabbit CRY1 | Proteintech | Cat# 13474-1-AP<br>RRID: AB_10697652 |
| Anti-Rabbit ERα | Cell Signaling Technology | Cat# 8644<br>RRID: AB_2617128 |
| Anti-Rabbit HER2 | Cell Signaling Technology | Cat# 2165<br>RRID: AB_10692490 |
| Anti-Rabbit Ki-67 | Abcam | Cat# ab16667<br>RRID: AB_302459 |
| Anti-Rabbit PER2 | Abclonal | Cat# a5107<br>RRID: 2863447 |
| Anti-Rabbit Vimentin | Abcam | Cat# ab92547<br>RRID: AB_10562134 |
| **Primers** |  |  |
| **Bacterial and virus strains** |  |  |
| NEB Stable Competent E. coli (High efficiency) | New England Biolabs | Cat# C3040H |
| **Biological samples** |  |  |
| Human breast samples | MCRC Biobank | Ref: 21_QIME_01 |
| TCGA-BRCA samples | FireBrowse | http://firebrowse.org/ |
| GTEX v8 breast samples | GTEX portal<sup>40</sup> | https://storage.googleapis.com/gtex_analysis_v8/rna_seq_data/GTEX_Analysis_2017-06-05_v8_RNASeQCv1.1.9_gene_tpm.gct.gz |
| BA11 brain samples | NIH GEO: GSE71620 | Chen et al.<sup>76</sup> |
| Groningen and Laval lung samples | NIH GEO: GSE23546 | Bossé et al.<sup>77</sup> |
| **Chemicals, peptides, and recombinant proteins** |  |  |
| 1X Trypsin-EDTA solution | Sigma-Aldrich | Cat# T3924 |
| 10X Tris/Glycine/SDS | Bio-Rad | Cat# 1610732 |
| A83-01 | Tocirs | Cat# 2939 |
| Acetic Acid Glacial | Fisher scientific | Cat# 10171460 |
| Advanced DMEM/F12 | Gibco | Cat# 12634010 |
| Agarose | Bioline | Cat# BIO-41025 |
| Alt-R S.p.Cas9 Nuclease V3 | Integrated DNA Technologies | Cat# 1081058 |
| Ampicillin | Sigma-Aldrich | Cat# A9393 |
| B27 supplement (50X), serum free | Gibco | Cat# 17504044 |
| BSA (Bovine Serum Albumins) | Sigma-Aldrich | Cat# A2153 |
| Calcium Chloride | Fisher scientific | Cat# 10171800 |
| Collagenase A | Roche | Cat# 10103186001 |
| Chloroform, 99.8+% | Fisher scientific | Cat# 10784143 |
| Dexamethasone | Sigma-Aldrich | Cat# D4902 |
| Dispase II | Roche | Cat# 37045800 |
| Donkey Serum | Sigma-Aldrich | Cat# D9663 |
| DPX Mountant for histology | Sigma-Aldrich | Cat# 06522 |
| DPBS (1X) | Sigma-Aldrich | Cat# D5652 |
| DMEM/F12 without phenol red | Gibco | Cat# 2063394 |
| Dnase I | Roche | Cat# 11284932001 |
| Dnase-free RNase set | QIAGEN | Cat# 79254 |
| Eosin-Y | Sigma-Aldrich | Cat# E4009 |
| EGF (Epidermal Growth Factor) | Peptotech | Cat# AF-100-15 |
| Ethanol, 99.8+% | Fisher scientific | Cat# 12337163 |
| FBS | Sigma-Aldrich | Cat# 12103C |
| FGF2 (Fibroblast Growth Factor 2) | Peptotech | Cat# 100-18B |
| FGF7 | Peptotech | Cat# 100-19 |
| FGF10 | Peptotech | Cat# 100-26 |
| Gentamicin solution | Sigma-Aldrich | Cat# G1272 |
| GlutaMax 100X | Gibco | Cat# 15630-056 |
| Goat Serum | Sigma-Aldrich | Cat# G9023 |
| Halt™ Protease inhibitor cocktails | Thermo Scientific | Cat# 78429 |
| HBSS | Sigma-Aldrich | Cat# H9394 |
| HCl (Hydrochloric Acid) | Fisher scientific | Cat# 10053023 |
| Hematoxylin Solution, Gill No. 2 | Sigma-Aldrich | Cat# GHS216 |
| HEPES | Sigma-Aldrich | Cat# H0887 |
| High-Capacity RNA to cDNA kit | Applied Biosystems | Cat# 4387406 |
| Hyaluronidases | Sigma-Aldrich | Cat# H3885 |
| Hydrogen peroxide solution | Sigma-Aldrich | Cat# 95321 |
| ISOLATE II Genomic DNA kit | Bioline | Cat# BIO-52066 |
| Isopropanol | Fisher scientific | Cat# 10477070 |
| L-Glutamine | Sigma-Aldrich | Cat# G7513 |
| Lipofectamine CRISPRMAX Transfection Regent | Thermo Scientific | Cat# CMAX00015 |
| Beetle Luciferin, Potassium Salt | Promega | Cat# E1603 |
| Lysing Matrix A | MP Biomedicals | Cat# 116910050 |
| Corning Matrigel® Growth Factor Reduced (GFR) Basement Membrane Matrix, LDEV-free | Corning | Cat# 354230 |
| Methanol | Fisher scientific | Cat# 10141720 |
| Methylcellulose | Sigma-Aldrich | Cat# M7027 |
| 4-20% Mini-PROTEIN TGX Precast Protein Gels | Bio-Rad | Cat# 4561093 |
| N-Acetylcysteine | Sigma-Aldrich | Cat# A9165 |
| Nicotinamide | Sigma-Aldrich | Cat# N6036 |
| Noggin | Peptotech | Cat# 120-10C |
| Opti-MEM™ I Reduced Serum Medium, no phenol red | Thermo Scientific | Cat# 11058021 |
| P/S (Penicillin/Streptomycin) | Sigma-Aldrich | Cat# P4333 |
| Polybrene | Sigma-Aldrich | Cat# TR-1003-G |
| Paraformaldehyde | Sigma-Aldrich | Cat# 95321 |
| Pierce™ BCA Protein Assay Reagent A | Thermo Scientific | Cat# 23223 |
| Pierce™ BCA Protein Assay Reagent B | Thermo Scientific | Cat# 23224 |
| Primocin | Invivogen | Cat# Ant-pm-1 |
| PureCol EZ gel solution | Sigma-Aldrich | Cat# 5074 |
| QIAfilter Plasmid Maxi Kit | QIAGEN | Cat# 12245 |
| QIAquick Gel Extraction Kit | QIAGEN | Cat# 28704 |
| Qubit™ RNA XR Assay Kit | Thermo Scientific | Cat# Q33224 |
| R-spondin 3 | Peprotech | Cat# 120-44 |
| RIPA Buffer | Sigma-Aldrich | Cat# R2078 |
| RNAlater™ Stabilization Solution | Thermo Scientific | Cat# AM7020 |
| RNA ScreenTape | Agilent | Cat# 5067-5576 |
| RNA ScreenTape Sample Buffer | Agilent | Cat# 5067-5577 |
| Rneasy Lipid Tissue Mini Kit | QIAGEN | Cat# 74804 |
| SB202190 | Sigma-Aldrich | Cat# S7067 |
| SIGMAFAST™ 3,3'-Diaminobenzidine tablets | Sigma-Aldrich | Cat# D4168 |
| Sodium Butyrate | Sigma-Aldrich | Cat# B5887 |
| Sodium citrate (100X) | Abcam | Cat# ab93678 |
| Streptavidin, Peroxidase (Concentrate, for IHC) | Vector | Cat# SA-5004-1 |
| TAE Buffer (50X) | Thermo Scientific | Cat# B49 |
| Takyon ROX probe Mastermix DTTP blue | Eurogenetic | Cat# EUK212 |
| Trans-Blot Tuobo RTA Mini 0.2 µm Nitrocellulose Transfer Kit, for 40 bolts | Bio-Rad | Cat# 1704270 |
| Triton X-100 | Thermo Scientific | Cat# 85111 |
| Tween™ 20 | Thermo Scientific | Cat# 85113 |
| Xylenes | Fisher scientific | Cat# 15618420 |
| Software and algorithms |  |  |
| STAR(version 2.5) | Dobin et al.<sup>78</sup> | https://github.com/alexdobin/STAR/releases |
| RSEM(version 1.3.0) | Li et al.<sup>79</sup> | https://github.com/deweylab/RSEM/releases |
| R (v4.2.1) | - | https://www.R-project.org |
| CCMapp | Wu et al.<sup>38</sup> | https://github.com/gangwug/CCMapp |
| BioRender | - | https://www.biorender.com |
| Julia (v1.6) | Bezanson et al.<sup>80</sup> | https://doi.org/10.1137/141000671 |
| GSEA (v4.2.3) | Subramanian et al.<sup>47</sup> | https://doi.org/10.1073/pnas.0506580102 |
| PSEA | Zhang et al.<sup>46</sup> | https://doi.org/10.1177/0748730416631895 |
| enrichR.R (v3.1) | Chen et al.<sup>48</sup> | https://doi.org/10.1093/nar/gkw377 |
| Dataframes.jl (v1.3.4) | - | https://dataframes.julidata.org/stable/ |
| Statistics.jl | - | https://docs.julialang.org/en/v1/stdlib/Statistics/ |
| StatsBase.jl (v0.33.21) | - | https://juliastats.org/StatsBase.jl/stable/ |
| LinearAlgebra.jl | - | https://docs.julialang.org/en/v1/stdlib/LinearAlgebra/ |
| MultivariateStats.jl (v0.10.0) | - | https://juliapackages.com/p/multivariatestats |
| Flux.jl (v0.13.5) | Innes.<sup>81</sup> | https://doi.org/10.21105/joss.00602 |
| PyPlot.jl (v2.11.0) | - | https://github.com/JuliaPlots/Plots.jl |
| Distributed.jl | - | https://docs.julialang.org/en/v1/stdlib/Distributed/# |
| Random.jl | - | https://docs.julialang.org/en/v1/stdlib/Random/ |
| CSV.jl (v0.10.4) | - | https://csv.juliadata.org/stable/ |
| Revise.jl (v3.4.0) | - | https://timholly.github.io/Revise.jl/stable/ |
| Dates.jl | - | https://docs.julialang.org/en/v1/stdlib/Dates/ |
| Multipletesting.jl (v0.5.1) | - | https://zenodo.org/badge/latestdoi/27935122 |
| sva.R (v3.44.0) | Leek et al.<sup>82</sup> | https://doi.org/doi:10.18129/B9.bioc.sva |
| refGenome (v1.7.7) | - | https://cran.r-project.org/src/contrib/Archive/refGenome/ |
| RTCGA.clinical (20151101.20.0) | - | https://bioconductor.org/packages/release/data/experiment/html/RTCGA.clinical.html |
| RTCGA (1.20.0) | - | https://www.bioconductor.org/packages/release/bioc/html/RTCGA.html |
| tidyverse.R (v1.3.2) | - | https://doi.org/10.21105/joss.01686 |
| aod.R (v1.3.2) | - | https://cran.r-project.org/package=aod |
| circular.R (v0.4-95) | - | https://r-forge.r-project.org/projects/circular/ |
| Other |  |  |

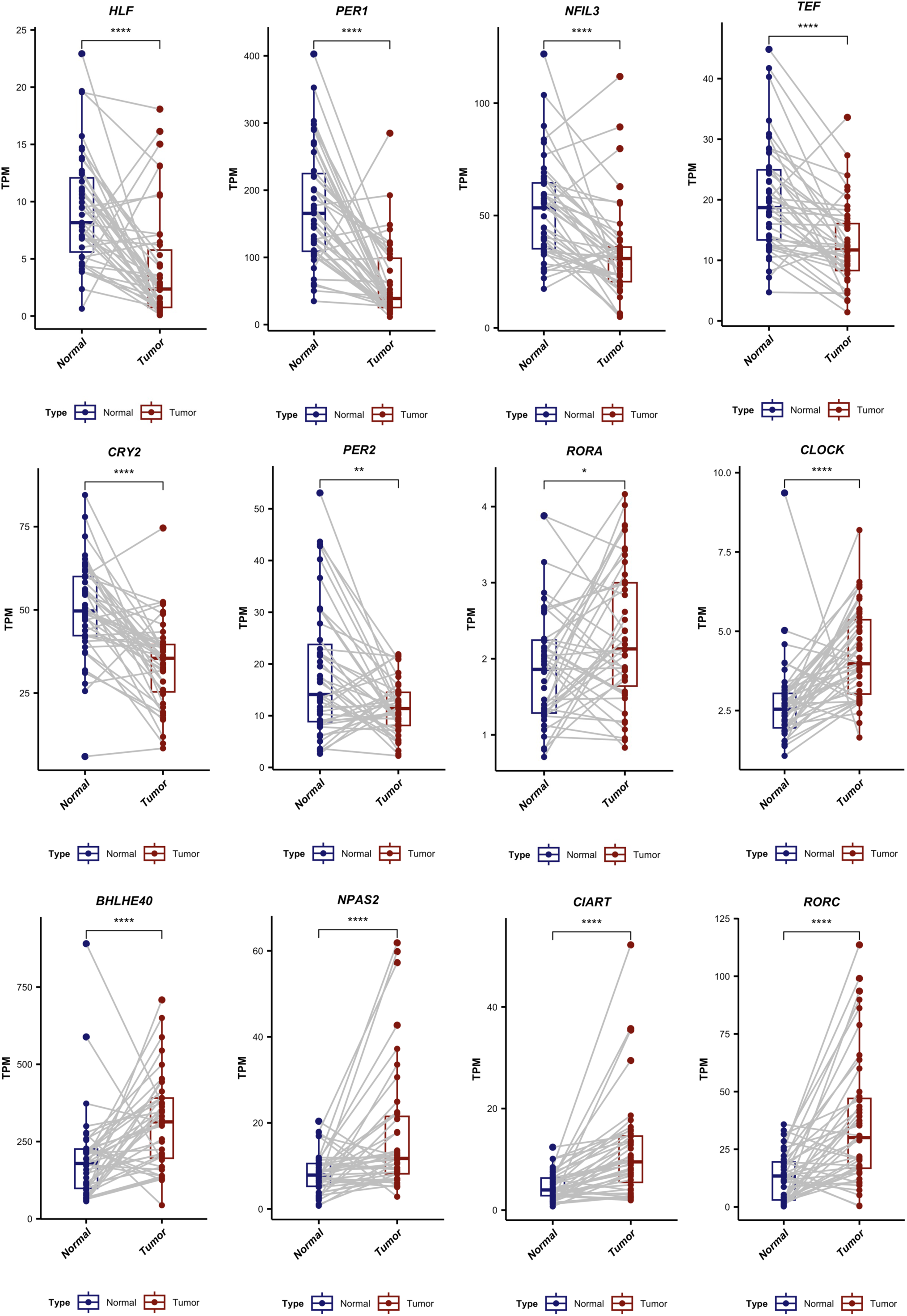
Fig S1.

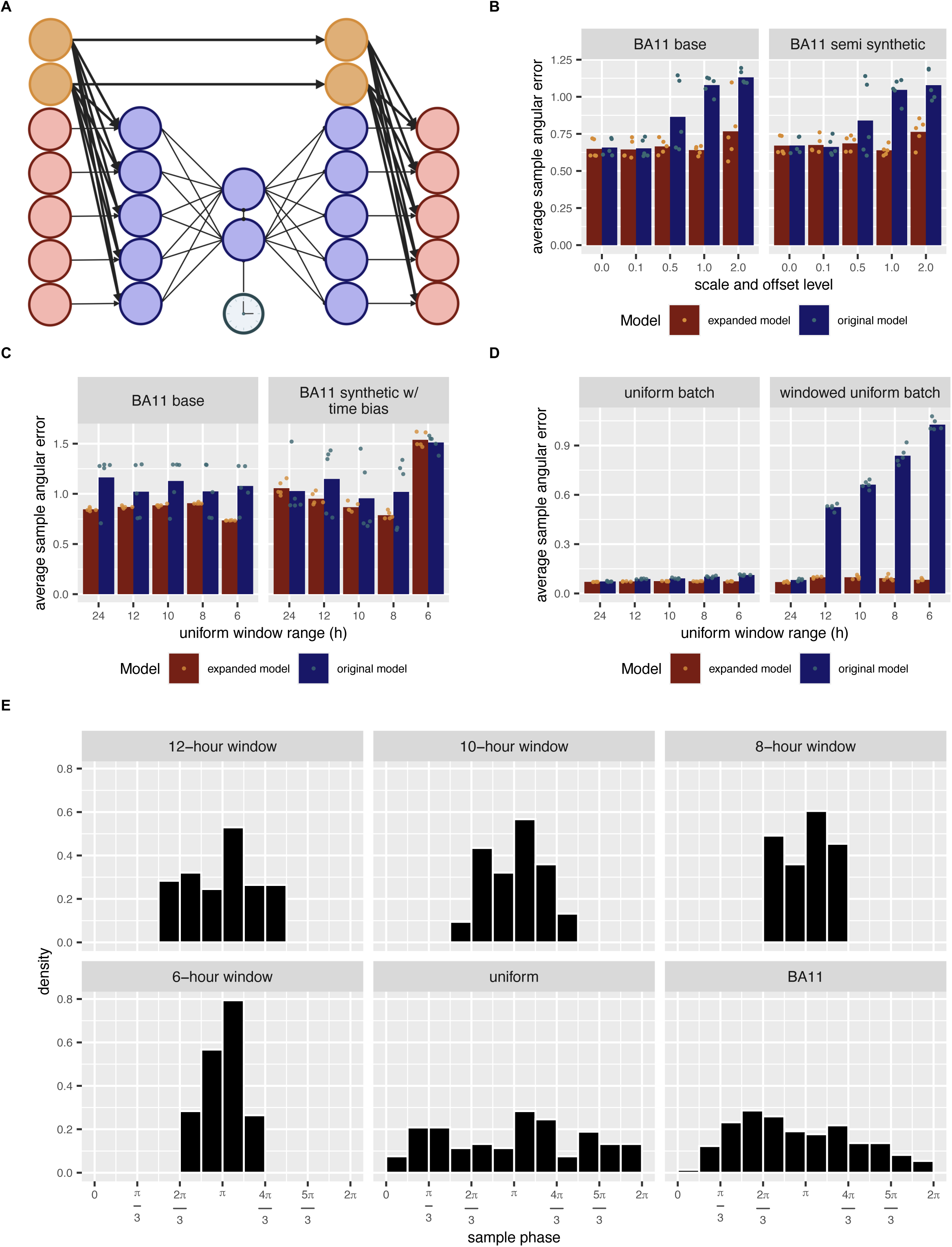
Fig S2.

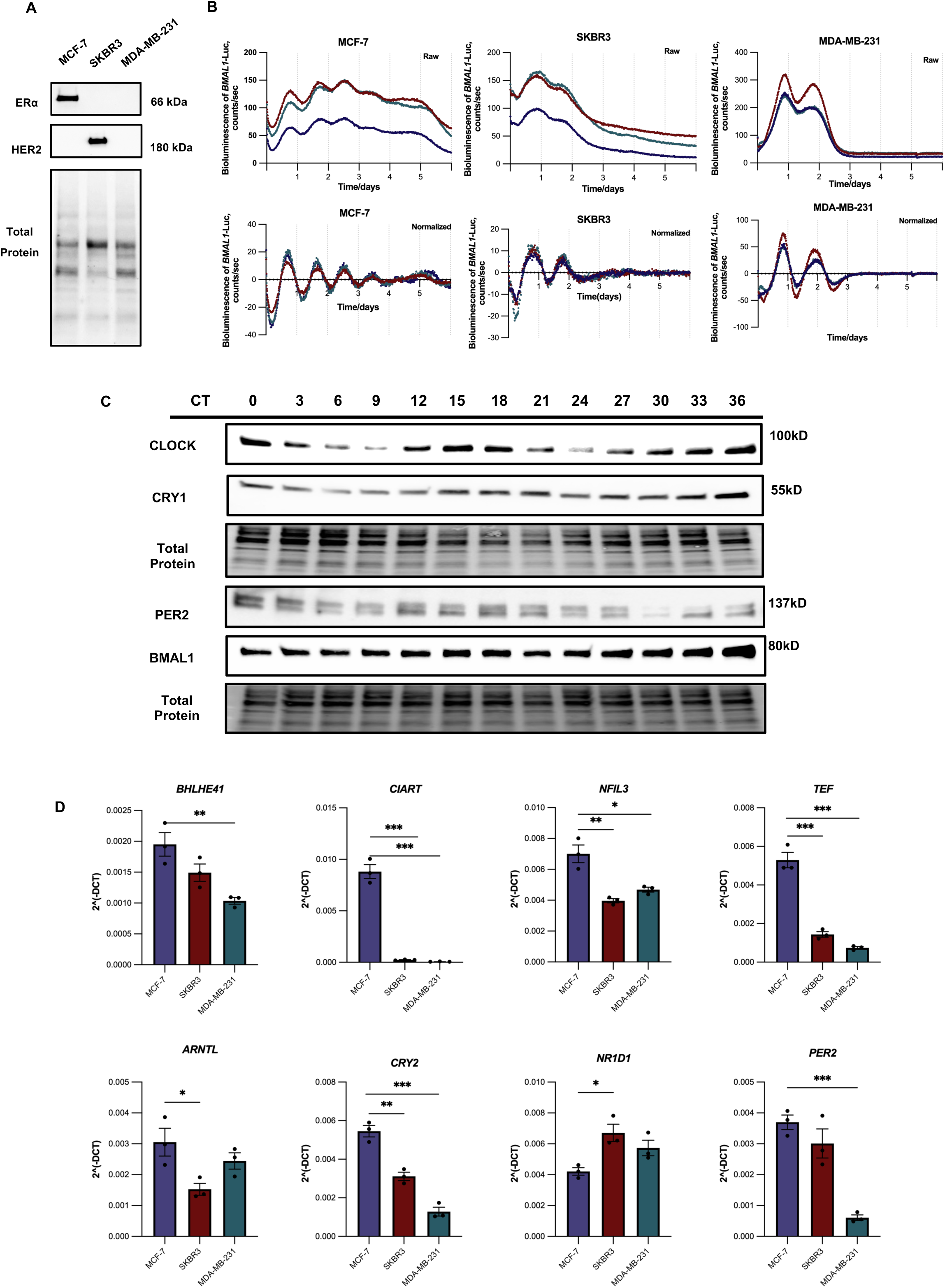
Fig S3.

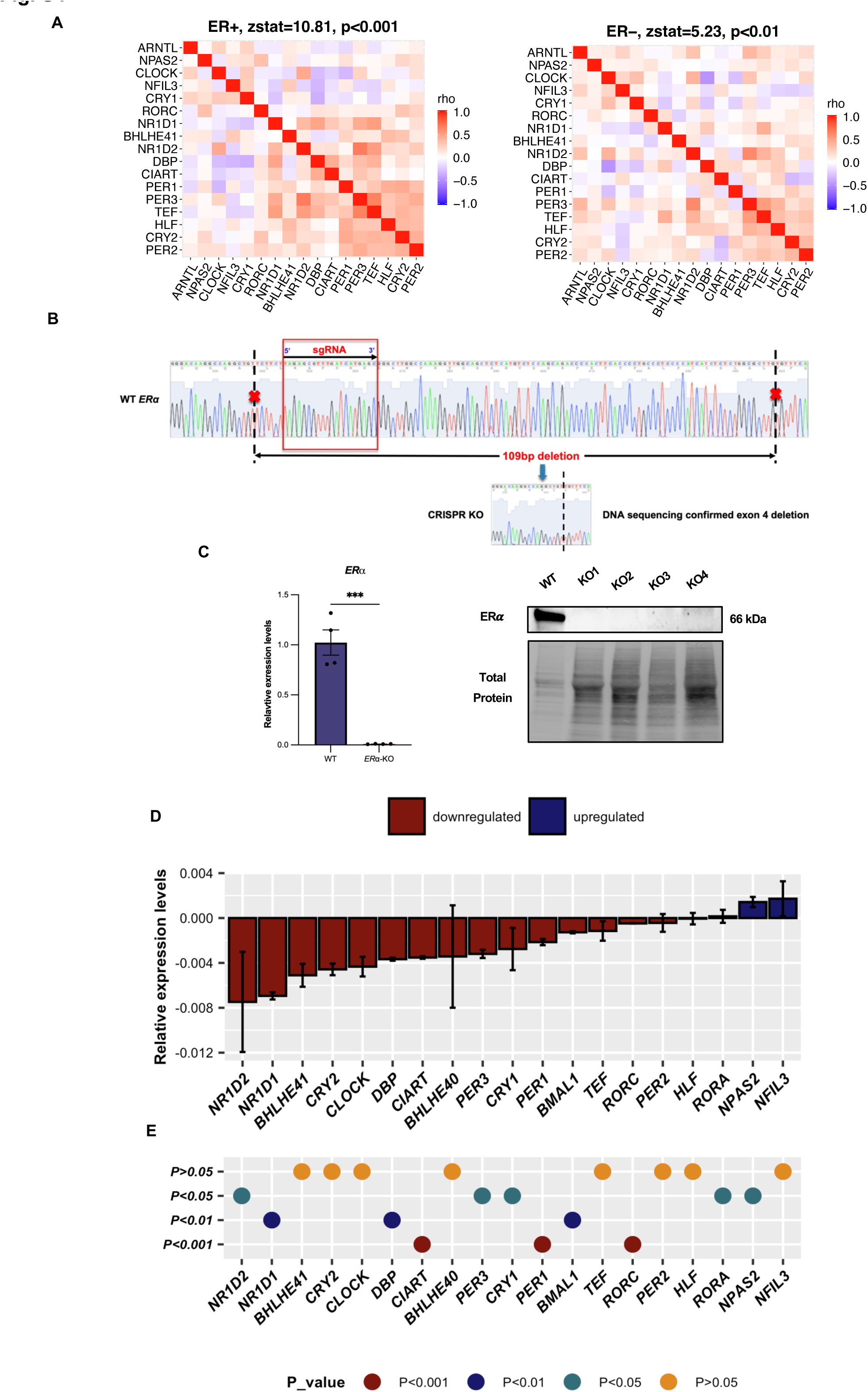
Fig S4.

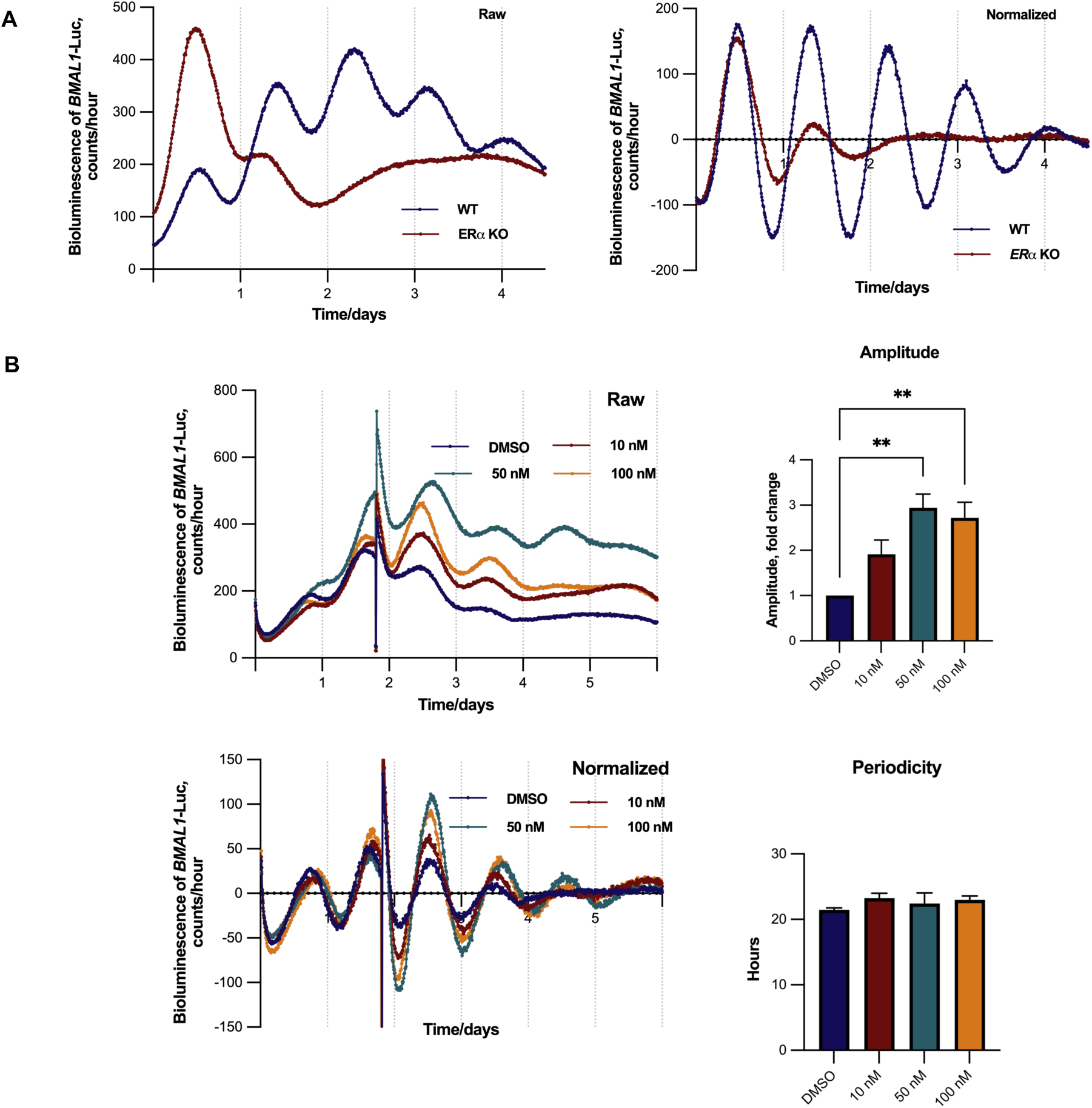
Fig S5.

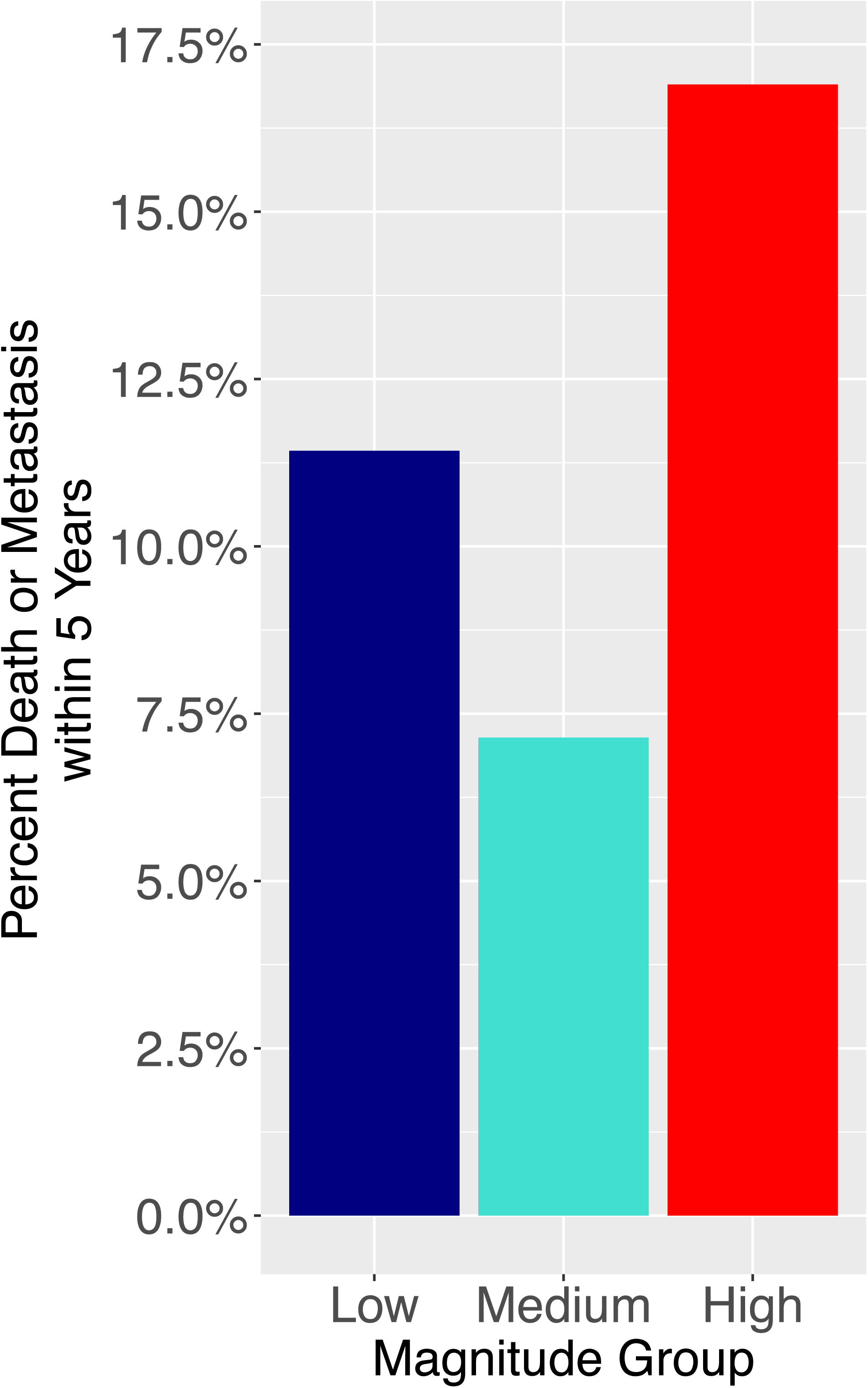
Fig S6.

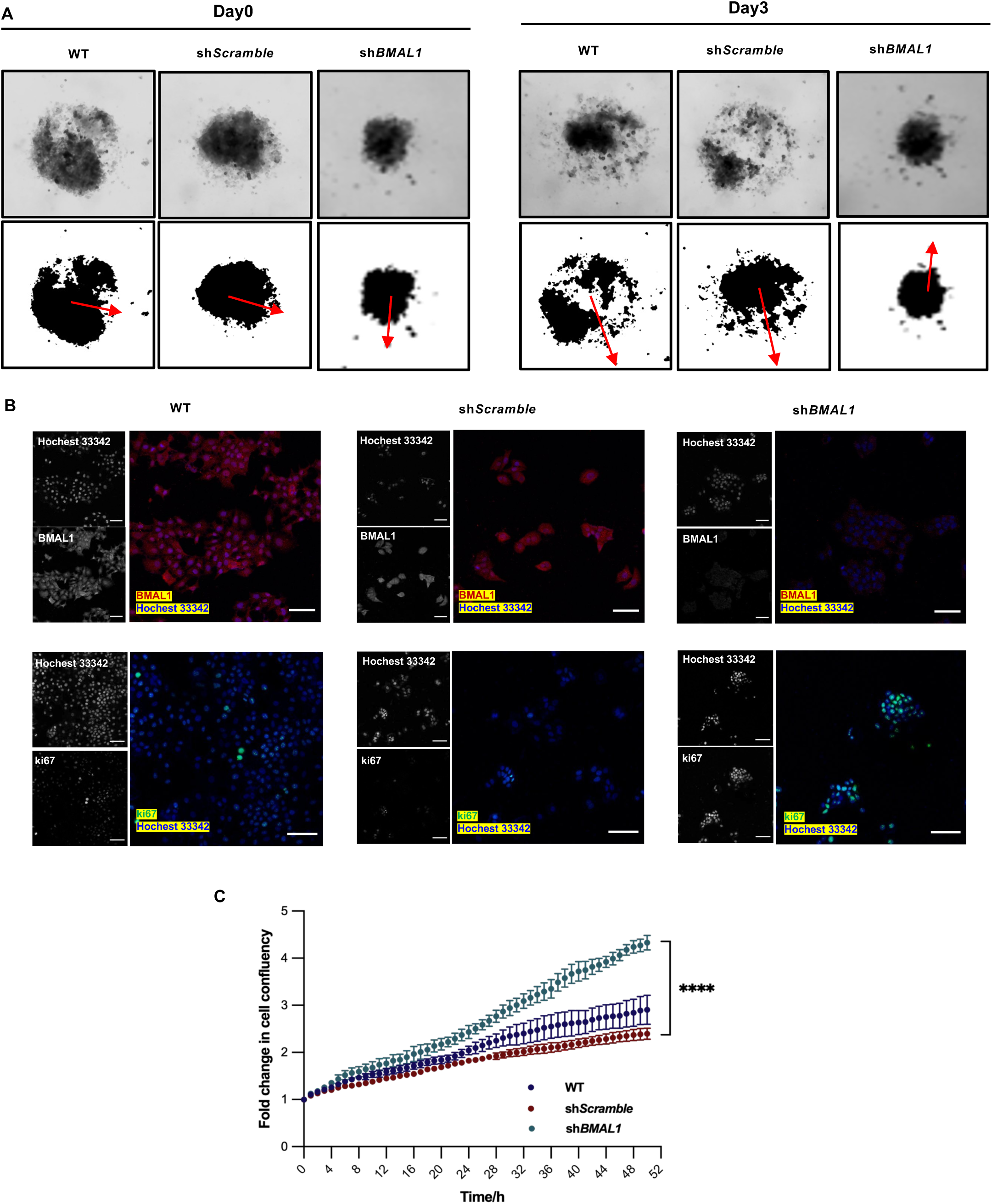
Fig S7.

