## Supplementary material for "Subtype-specific circadian clock dysregulation modulates breast cancer biology, invasiveness, and prognosis": Figure Titles and Captions

**Figure titles and legends**

**Fig. 1 RNAseq of paired clinical samples revealed widespread clock changes in breast tumors**

(A) A diagram of our hybrid study design characterizing circadian clocks in non-cancerous and cancerous human breast tissues: We integrated time-stamped biopsies with existing breast cancer databases and used informatic ordering to reconstruct rhythms. (1) The Manchester Cancer Research Centre Biobank provided breast tissue samples along with documentation of resection time, histologic diagnosis, and patient demographics. Matched, non-cancerous samples were obtained at least 4 cm away from tumor samples. After histologic verification (2), primary cells were isolated and bulk tissue was sent for RNAseq analysis. (3) Core clock gene correlation pattens were assessed in different tumor subtypes using public data. A bioluminescent *BMAL1*-Luc circadian reporter was used to evaluate rhythms in different cancer subtypes using human primary tumor organoids. (4) CYCLic Ordering by Periodic Structure (CYCLOPS) was used to reconstruct rhythms in non-cancerous breast tissue and in Luminal A cancers. The influence of circadian clocks on patient prognosis and cancer cell behavior were further determined by magnitude analysis and experimental studies.
(B) A pie chart depicts the number of each cancer subtype included in the 43 pairs of time-stamped clinical samples: Luminal A (29 pairs), Luminal B (3 pairs), HER2 (9 pairs) and Triple Negative Breast Cancer (2 pairs).
(C) A rose plot depicts the resection time of the 43 Manchester samples. Dark blue and dark red indicate samples collected in the morning (9:00 – 12:00) or afternoon (12:00 – 17:00), respectively.
(D) Representative Hematoxylin & Eosin-Y (H&E) staining and immunohistochemistry validating normal and malignant biopsies in paired human breast tissues. Paired samples were immunostained with Cytokeratin 8 (CK8, an epithelium marker) and Vimentin (a stromal marker). Nuclei were counterstained with hematoxylin. Scale bar= 100 µm, N=43
(E) Top: Bar chart showing log_2_ fold changes in the mRNA expression levels of core clock genes in human breast tumors as compared to matched normal samples, N=43. Bottom: Dot plots displaying *p*-values assessing differential expression, N = 43, pairwise Wilcoxon test (non-parametric).
(F) Evaluation of core circadian organization in human breast tissue. Heatmaps of Spearman’s ρ for clock and clock-associated genes from non-cancerous breast and tumor samples with a sequencing depth of > 20 million reads are shown. The zstat value and p-value are computed using a Mantel test and a reference correlation matrix of clock and clock-associated genes from the mouse atlas data. Higher zstat scores denote a stronger resemblance to the established reference for healthy tissues.

**Fig. 2 CYCLOPS reconstructed rhythms in non-cancerous human breast samples**

CYCLOPS was used to estimate sample circadian phase in non-cancerous human breast tissue. Some samples were newly collected in Manchester, UK (N = 26). Most data were obtained from GTEx (N = 167) and TCGA (N =106).

(A) Inferred time of peak transcript expression phase (acrophase) is plotted for select core clock genes in non-cancerous human breast (outer, blue). The acrophases for the mouse paralogues (averaged across mouse tissues) are shown in the inner circle (inner, orange). In the mouse, time 0 was defined by the peak time of *Arntl* (*Bmal1*) expression. Rhythmicity for the transcripts shown was assessed by modified cosinor regression. All transcripts in blue had a regression p-statistic < 0.05. The Human ordering was aligned to match the mouse acrophases.
(B) Histogram of inferred sample collection phase for non-cancerous GTEx (filled) and TCGA (outlined) data. Sample phases are aligned as in (A).
(C) Optimal alignment of CYCLOPS-predicted sample phases for the subset of time-stamped samples collected in Manchester (${Corr}_{Fisher}=0.69, p_{Fisher}<0.01$).
(D) Histogram of transcript acrophases including all significantly cycling transcripts in non-cancerous breast tissue (BHq < 0.05).
(E) Transcript expression was fit using cosinor regression in a model that included batch collection site. Batch-adjusted transcript expression is plotted as a function of CYCLOPS-predicted sample phase. The best-fit sinusoid is superimposed, and fit significance is shown for each transcript. Fit significance is assessed by modified cosinor regression. Solid lines represent a significant cosinor fit (BHq < 0.05).
(F) Phase set enrichment analysis (PSEA) was applied to CYCLOPS ordered non-cancerous human breast data. MSigDB hallmark gene sets exhibiting phase-coordinated expression are shown. Gene sets graphed furthest from the center of the circle had the most significant phase coordination. Gene sets highlighted in yellow also demonstrated phase coordination in mouse mammary tissue.
(G) EnrichR was used to identify MSigDB Hallmark gene sets overrepresented among informatically identified cycling genes in human non-cancerous breast (BHq < 0.05, relative amplitude > 0.33). The significance of pathway overrepresentation (log BHq) is plotted against the significance of pathway phase coordination identified by PSEA (log BHq).
(H) Transcripts were ranked by cosinor regression F statistic. Gene set enrichment analysis (GSEA) was applied to the ranked list. Hallmark gene sets with BHq < 0.1 are shown.

**Fig. 3 Subtype specific circadian rhythm dysfunction in breast cancers**

(A) Evaluation of core circadian organization in human breast cancer subtypes from TCGA data. Heatmaps depicting the Spearman correlation between selected core clock genes are shown for non-cancerous breast tissue (N=111), Luminal A (N=532), Luminal B (N=203) and Basal (N=181). The zstat value and p-value are computed using a Mantel test and a reference correlation matrix of clock and clock-associated genes from the mouse atlas data. Higher zstat scores denote a stronger resemblance to the established reference for healthy tissues.
(B) Left, Representative images of mammary organoids derived from non-cancerous breast and paired tumor tissues from the same individual, scale bar=50 µm. Right, Immunostaining demonstrates morphological structures of non-cancerous mammary organoids and matched tumor organoids. Hoechst 33342 was used for nucleus and F-Actin was used for cytoskeleton, scale bar=50 µm.
(C) Circadian rhythms in human normal mammary organoids and matched tumor organoids were monitored by a LV200 bioluminescence imaging system. Representative bioluminescence images of organoids transfected with *BMAL1*-Luc from Luminal A (N=4) and TNBC (N=3) subtypes were taken at 6-hour intervals (CT15, CT21, CT27, CT33 and CT39), scale bar=100 µm. Both raw and detrended bioluminescence signals are shown. (dark red traces, Luminal A organoids; dark blue traces, non tumor organoids). Rhythmicity was assessed by Olympus LV200 bioluminescence imaging system (dark red traces, normal organoids; dark blue traces, tumor organoids). Notably, non-cancerous and tumor organoids were recorded in separate microscope runs and bioluminescence images from tumor organoids were adjusted for ease of visualization.

**Fig. 4 CYCLOPS reconstructed circadian rhythms from Luminal A samples**

CYCLOPS was used to estimate sample circadian phase in Luminal A samples. These data included n = 18 samples from Manchester and n = 193 samples from TCGA.

(A) Inferred acrophase are plotted for select core clock genes in Luminal A human breast tumors (outer, blue) and for the mouse paralogues (inner, orange). Mouse acrophases were determined from the circular average across tissues in the Mouse Circadian Atlas. In the mouse, time 0 is defined by the circular average peak time of *Arntl* expression. Rhythmicity for the transcripts shown was assessed by modified cosinor regression. All transcripts in blue had a regression p-statistic < 0.05. The Human ordering was aligned to match the mouse acrophases.
(B) Transcript expression in Luminal A samples was fit by cosinor regression in a model that included batch collection site. Separate models were fit to Luminal A and non-cancerous samples. Batch-adjusted transcript expression is plotted as a function of CYCLOPS-predicted sample phase. Luminal A samples are shown in red and non-cancerous samples are shown in blue. The best-fit sinusoid is shown for each condition. Solid lines represent a significant cosinor fit (BHq < 0.05), and dashed lines represent a non-significant fit.
(C) Histogram of transcript acrophases including all significant cycling transcripts (BHq < 0.05) identified in Luminal A breast tissue, by modified cosinor regression.
(D) A histogram shows the log amplitude ratio comparing Luminal A and non-cancerous samples. Transcripts that cycled in either Luminal A or non-cancerous samples (BHq < 0.05) were included.
(E) The CYCLOPS estimated phase of Luminal A tumor samples is plotted against the matched non-cancerous samples for the small number of matched samples in TCGA. Little correlation is evident.
(F) MSigDB hallmark gene sets exhibiting phase-coordinated expression are shown. PSEA was applied to CYCLOPS-ordered Luminal A data. Gene sets graphed furthest from the circle's center had the most significant phase coordination. All gene sets other than those highlighted in red also demonstrated phase coordination in non-cancerous breast samples.

**(G)&(I)** MSigDB Hallmark gene sets enriched for increased and decreased cycling in Luminal A samples are shown in G & I, respectively. Transcripts significantly cycling in either Luminal A or non-cancerous samples (BHq < 0.05) were ranked by the log fold amplitude change and analyzed by GSEA.

**(H)&(J)** MSigDB Hallmark gene sets overrepresented among transcripts with a 5-fold increase or 10-fold decrease in Luminal A sample amplitude are shown in H & J, respectively. Circadian transcripts that showed significantly differential cycling in Luminal A and non-cancerous tissues were identified in a nested model. The base model included the collection site and Luminal A/non-cancerous specific mean expression. The expanded model added Luminal A/non-cancerous specific cycling parameters.

**Fig 5 CYCLOPS clock magnitude varies in Luminal A samples, correlates with EMT, and predicts prognosis.**

(A) Cartoon depicting CYCLOPS sample magnitude. CYLOPS projects the n-dimensional eigengene expression of each sample to a plane where the circular data structure is apparent. The angular position of any sample in the reconstructed circle reflects its circadian phase. The distance from the circle's center to a given sample is the sample’s CYCLOPS magnitude (dotted line). This distance is a weighted sum of the amplitudes of individually cycling seed genes.
(B) Histogram of Luminal A CYCLOPS sample magnitudes colored by tertiles (N = 70). The distribution shows a long tail.
(C) A histogram of transcript amplitude ratios comparing samples from the top-third and bottom-third magnitude groups. Nested cosinor regression models assessed the significance of including CYCLOPS sample magnitude in fitting transcript rhythms. Transcripts that showed differential cycling by CYCLOPS magnitude group were identified (BHq <0.05). For differentially cycling transcripts the amplitude of cycling in the highest tertile was compared to the amplitude of cycling in the lowest tertile.
(D) Batch normalized expression data are shown for representative core clock transcripts that exhibit differential cycling between magnitude groups (BHq <0.05). Luminal A samples in the top tertile of CYCLOPS magnitude are shown in red and the bottom two tertiles in blue. The best-fit sinusoid is superimposed for each group. Significance was assessed by nested cosinor regression.
(E) The CYCLOPS estimated phase of Luminal A tumor samples is plotted against the matched non-cancerous samples for the small number of matched samples in TCGA. Samples are colored by CYCLOPS magnitude group, distinguishing the top tertile from the bottom two tertiles.
(F) Five-year patient mortality grouped by tumor CYCLOPS magnitude group. Patient outcome data were obtained from the TCGA database. For each tertile/magnitude group, the percentage of patients who died within 5 years of diagnosis is shown. The increased risk in the high-magnitude tumor group remained statistically significant when evaluated in a logistic regression model that included patient age and the presence of known metastasis at diagnosis (chisq p<0.05).
(G) MSigDB Hallmark gene sets overrepresented among transcripts that showed differential cycling among Luminal A magnitude groups are shown. Transcripts showing differential cycling in Luminal A magnitude groups (BHq < 0.05) were input to EnrichR, and overrepresented gene sets were identified. Similarly, all transcripts cycling in Luminal A tumors were ranked by the ratio of transcript amplitude in the highest tertile compared to the lowest. GSEA was used to identify MSigDB Hallmark gene sets enriched in the ranked list. The -log(BHq) from the EnrichR analysis is plotted against the normalized enrichment score (NES) from GSEA. Both methods identify the “Epithelial-mesenchymal transition” gene set as most impacted by differential cycling in the high magnitude group.
(H) Batch normalized expression data for selected transcripts in the Epithelial-mesenchymal transition pathway exhibit differential cycling (BHq <0.05). Luminal A samples in the top tertile of CYCLOPS magnitude are shown in red and the bottom two tertiles in blue. The best-fit sinusoid is superimposed for each condition.

**Fig. 6 Invasion of MCF-7 cells into 3D collagen matrix is suppressed by circadian clock disruption.**

(A) Representative traces of circadian rhythms in *BMAL1*-Luc expression in MCF-7 cells transduced with lentiviral sh*Scramble* or sh*BMAL1*. Non-transduced cells (WT) were also included as an additional control. Left, raw data; right, normalized data. N = 3.
(B) MCF-7 cell invasion was assessed by a hanging drop cell invasion assay in a 3D collagen matrix. Representative images of cell migration are displayed at Day 0 (left) and Day 3 (right). Mask option on ImageJ was used to convert original images to 8-bit images, aiding visualization and quantification of distance of cell migration.
(C) The relative distance of cell migration was quantified and plotted. Data were normalized to WT which was set as 1. Unpaired t-Test, ^**^*p*< 0.01, N=4.
(D) Representative traces of circadian rhythms in *BMAL1*-Luc expression in MCF-7 cells treated with different doses of KL001 (CRY1/CRY2 stabilizing compound).
(E) Representative images of cell invasion assay of MCF-7 cells treated with KL001.
(F) Quantification of cell migration distance in (E), Unpaired t-Test, ***p*<0.01 N = 3.

**Supplemental Figure Legends and Titles**

**Supplementary 1, Fig. S1-S7 Table. S1-S2**

**Fig. S1 Differential mRNA expression levels of core circadian clock genes in human breast tumors as compared to paired normal tissues (Related to Fig. 1)**

N = 43, pairwise Wilcoxon test (non-parametric), *^*^p*<0.05, *^**^p*<0.01, *^****^p*<0.0001.

**Fig. S2 CYCLOPS 2.0 outperforms CYCLOPS in benchmarking with confounded and time-biased data (Related to Fig. 2)**

CYCLOPS 2.0 was benchmarked on semi-synthetic and synthetic data to test the influence of confounding batch effects and biases in sample collection times on the ability to order rhythmic data.

(A) Depiction of CYCLOPS 2.0, an autoencoder algorithm with built-in batch/covariate normalizing layers. The blue circles represent the original architecture of CYCLOPS. The red and the yellow circles represent the *covariate correction wrapper*, the CYCLOPS 2.0 expansion. The clock face in the center, at the bottom, represents any sample collection times that may be provided to guide the ordering process.
(B) Benchmarking of CYCLOPS and CYCLOPS 2.0 on semi-synthetic BA11 data at various levels of confounding batch influence. We used an autopsy-based collection of BA11 cortical human gene expression (Chen et al, 2016) with well annotated collection time as base data. We then used the observed differences between two large clinical lung biopsy datasets (Bossé et al.^1^) to model realistic differences in expression expected from different collection and processing sites. We generated BA11 expression data from an idealized second center. Using the differences in scale and offset observed in the clinical pulmonary datasets as a baseline, we generated synthetic datasets with more pronounced or less pronounced differences in expression. At each level of difference, five semi-synthetic datasets were generated. Scale and offset differences observed between the two lung datasets are termed level 1.0. More and less extreme differences are obtained by exponentiating the modeled log normal distributions. Levels less than 1 reflect more modest differences than those observed in the lung data sets. Levels greater than 1 reflected more marked differences. Both the original CYCLOPS and CYCLOPS 2.0 were used to time order these data. The average sample angular error was calculated within each batch, for each of the best models. Each batch contained 202 samples, for 404 total samples in each semi-synthetic dataset.
(C) Benchmarking of CYCLOPS and CYCLOPS 2.0 on synthetic BA11 data that included both batch influence and differential bias in sample collection time. The original BA11 dataset included samples from across the circadian day with little temporal bias. Using the observed transcript cycling parameters we created a fully synthetic BA11 dataset that included batch confounding (as above) but only included samples from a limited temporal window. This modeled a clinical collection. The sample collection times of the synthetic BA11 batch were drawn from random uniform distributions with ranges of 24, 12, 10, 8, or 6 hours. In each case the scale and offset levels were set to 1, modeling the differences observed in the two lung datasets. For each temporal window, five synthetic datasets were generated. The original CYCLOPS was applied after ComBat normalization. CYCLOPS 2.0 was applied without ComBat normalization but including batch identifiers as a covariate. The average sample angular error was calculated within each batch, for each of the best models. The synthetic data included half as many samples as the original data.
(D) Benchmarking of CYCLOPS and CYCLOPS 2.0 on synthetic pulmonary data that included both batch influence and differential bias in sample collection time. We repeated the above analysis using data from lung tissue. Lung tissue has been shown to have many more cycling transcripts. We used CYCLOPS to separately order the Laval and Groningen pulmonary datasets in Bosse et al.^1^ as we had previously done (Anafi et al, 2017). We then estimated the cycling parameters for all cycling transcripts and again modeled the observed differences between the Laval and Groningen datasets. We created data from two idealized centers. A synthetic uniform batch (covers the entire 24-hour period) that mimics the expression parameters in the Groningen data was created. A synthetic dataset that mimics the expression parameters of the Laval data was separately created. The sample collection times in the synthetic “Laval” batch were drawn from random uniform distributions with ranges of 24, 12, 10, 8, or 6 hours. For each temporal collection range, five synthetic datasets were generated. Data were ComBat normalized prior to application of the original CYCLOPS. CYCLOPS 2.0 was applied without ComBat normalization but including batch identifiers. The average sample angular error was calculated within each batch, for each of the best models.
(E) Representative histograms of simulated sample collection times and the observed collection times of the BA11 (Chen et al.^2^) data.

**Fig. S3 Circadian rhythms in different breast cancer cell lines (Related to Fig. 3)**

(A) Representative Western Blots showing protein expression levels of ERα and HER2 in different subtypes of breast cancer cell lines (MCF-7, SKBR3 and MDA-MB-231). Total protein levels were used for normalization. N=3.
(B) Bioluminescence recordings of *BMAL1*-Luc levels in three different breast cancer cell lines. N=3 replicates are shown for each cell line.
(C) Representative Western Blots of endogenous levels of CLOCK, CRY1, PER2 and BMAL1 in MCF-7 cells over 36 hours at 3-hour intervals. Cells were synchronized with dexamethasone. CT0 is defined as 24 hours after synchronization. Total protein levels were used for normalization. N=3.
(D) Core clock mRNA expression levels are shown for three breast cancer cells lines: MCF-7, SKBR3 and MDA-MB-231. Cells were not synchronized so as to assess the average transcript levels. ^*^*p<*0.05, ^**^*p<*0.01, ^***^*p<*0.001, unpaired t-Test, N=3.

**Fig. S4 ERα deficiency disrupted molecular clocks in breast cancer cells (Related to Fig. 3)**

(A) Evaluation of core circadian organization in ER positive vs. ER negative human breast tumors in the TCGA database. Heatmaps of Spearman’s ρ for clock and clock-associated genes are shown. The zstat value and p-value are computed using a Mantel test and a reference correlation matrix of clock and clock-associated genes from the mouse atlas. Higher zstat scores denote a stronger resemblance to the established reference for healthy tissues.
(B) ERα-KO in MCF-7 cells using CRISPR-Cas9. sgRNA was designed to target exon 4 of *ERα* transcripts. A representative DNA sequencing confirmed gene targeting of exon 4.
(C) Validation of ERα deletion from MCF-7 cells. Left, qPCR for mRNA levels of *ERα* in WT and ERα-KO MCF-7 cells, unpaired t-Test, ^***^*p*<0.001, N=4. Right, representative Western Blot showing expression levels of ERα in MCF-7 cells with and without ERα-KO, N=4.
(D) Bar chart showing changes in mRNA expression levels of circadian clock genes in MCF-7 cells with ERα-KO vs. WT cells measured by qPCR, N=4.
(E) Dot plots displaying *p*-values assessing differential expression, N = 4, unpaired T-test.

**Fig. S5 ERα is required for robust circadian rhythms in MCF-7 breast cancer cells (Related to Fig. 3)**

(A) Representative bioluminescence traces of *BMAL1*-Luc reporter in WT and ERα-KO MCF-7 cells, N=4 individual KO clones.
(B) Left, Bioluminescence traces of *BMAL1*-Luc reporter in MCF-7 cells treated with an ERα selective agonist …(PPT) at various concentrations (10 nM, 50 nM and 100 nM). Cells were unsynchronized at the beginning of the experiment. Dimethyl Sulfoxide(DMSO) was used as control. Right, quantification of amplitude and period of oscillations following PPT treatment. One-Way ANOVA, ***p*< 0.01 vs. DMSO group, N=3.

**Fig. S6 Patient Outcome Analysis (Related to Fig. 5)**

Five-year patient mortality or new tumor event grouped by tumor CYCLOPS magnitude group. Patient outcome data were obtained from the TCGA database. For each tertile/magnitude group, the percentage of patients who died or had a new tumor event within 5 years is shown. The increased risk in the high-magnitude tumor group was not statistically significant when evaluated in a logistic regression model that included patient age and the presence of known metastasis at diagnosis (chisq *p*=0.10).

**Fig. S7 Molecular clock disruption in breast cancer cells compromises invasiveness but promotes cell proliferation (Related to Fig. 6)**

(A) Primary Luminal A cell invasion was measured by hanging drop cell invasion assay in a 3D collagen matrix. Representative images of cell migration are shown at Day 0 (left) and Day 3 (right). Mask option on ImageJ was used to convert original images to 8-bit images, aiding visualization and quantification of distance of cell migration. Red arrows show the furthest migration in each experiment.
(B) Immunofluorescent staining of BMAL1 and Ki-67 in MCF-7 cells lentivirally transduced with sh*Scramble* or sh*BMAL1*. Hoechst 33342 was used as a nuclear counterstain. N=3
(C) IncuCyte imaging was used to measure cell growth of MCF-7 cells following sh*BMAL1* knockdown, ^****^*p*< 0.0001, One-way ANOVA, N=3.

1.	Bossé, Y., Postma, D.S., Sin, D.D., Lamontagne, M., Couture, C., Gaudreault, N., Joubert, P., Wong, V., Elliott, M., van den Berge, M., et al. (2012). Molecular signature of smoking in human lung tissues. Cancer Res *72*, 3753-3763. 10.1158/0008-5472.Can-12-1160.

2.	Chen, C.-Y., Logan, R.W., Ma, T., Lewis, D.A., Tseng, G.C., Sibille, E., and McClung, C.A. (2016). Effects of aging on circadian patterns of gene expression in the human prefrontal cortex. Proceedings of the National Academy of Sciences *113*, 206-211. doi:10.1073/pnas.1508249112.
