## Supplementary material for "Subtype-specific circadian clock dysregulation modulates breast cancer biology, invasiveness, and prognosis": Methods

**Resource availability**

**Lead contact**

Further information and requests for resources and reagents should be directed to and will be fulfilled by the joint senior corresponding authors: Ron C Anafi and Qing-Jun Meng.

**Materials availability**

This study did not generate new unique reagents.

### Experimental model and subject details

Patient samples

All work involving fresh human tissues was approved by the Human Tissue Authority and the Ethical Approval Committee of the MCRC (Manchester Cancer Research Centre) Biobank at The Christie NHS Foundation Trust (Ref: 21_QIME_01). Tumor and paired normal tissues at least 4 cm away from the same breast of patients (Fig. 1A). All patients recruited from the Nightingale Breast Screening Clinic consented to use their tissues for biological research. Tissues dissected from the breast following mastectomy were dipped into DMEM/F-12 (Dulbeccon’s modified Eagle’s medium/Nutrient Mixture F-12 Ham Medium, Sigma-Aldrich) and sent to the laboratory at room temperature and kept at 4 °C in the lab till further tissue preparation. The tissues for RNA isolation were put in RNA*later* (Invitrogen) solution for room temperature transport. 43 pairs of fresh human samples were obtained to isolate RNA and conduct bulk RNA-seq, while 24 pairs were collected to isolate normal mammary epithelial cells (MECs) and breast tumor cells and to track circadian rhythms in distinct subtypes of breast cancer. The demographic information of breast cancer patients was documented on Table. 1, including age, gender, time of resection, receptor status, stage, grade and Ki-67.

Human cell lines

Cell lines including MCF-7, SKBR3 and MDA-MB-231 were kindly gifted by the Tournier Lab in the University of Manchester. MCF-7/HER2-18 cells^1^ were obtained from the Clarke Lab in the University of Manchester. HEK-293T cells were gifted by the Kimber Lab. All cells were cultured in DMEM (Sigma-Aldrich) with 10% FBS (Fetal Bovine Serum, Invitorgen), 1% L-Glutamine (Sigma-Aldrich) and 1% penicillin/streptomycin (Sigma-Aldrich).

### Method details

Isolation of primary MECs from human breast tissues

Human breast tissues were minced using scalpels and scissors and were then moved into 50 mL Falcon tubes with 12 mL DMEM/F12 media, 10% FBS, 1% L-Glutamine, 1% penicillin/streptomycin, 2.5 mg/mL Collagenase A solution (Roche), 1 mg/mL Hyaluronidase solution (Sigma-Aldrich). The samples were digested in a shaking incubator at 37 °C and 160 rpm overnight. Another day, MECs were isolated from human breast tissues using different centrifuge speed^2^. Digested tissues were centrifuged at 40 g for 1 min to isolate explants enriched MECs. The fat layer on the top of the supernatant was discarded, and the supernatant was transferred into a new Falcon to centrifuge at 100 g for 2 min to enrich more MECs. The supernatant was removed from Falcons and the two pellets were combined into one Flacon tube using HBSS (Sigma-Aldrich), then centrifuged again at 100 g for 2 min. Single cells were released from pellets in 3-5 mL of 1X Trypsin-EDTA (Sigma-Aldrich) with constant agitation using tips or pipettes. HBSS with 4% FBS was added to neutralize the trypsin, and the suspension was centrifuged at 40 g for 1 min. Pellets were suspended by HBSS to wash out the serum and were centrifuged at 40 g for 1 min. Later, pellets were resuspended in 4 mL HBSS with 5 mg/mL Dispase II (Roche) and 10 mg/mL DNase I (Roche) to remove DNA, and the washing steps were repeated. After centrifuging, MECs were further enriched in the pellets.

Lentiviral packaging

For lentivirus packaging with *Bmal1*-Luc/GFP, 12 μg *Bmal1-Luc*/GFP plasmids, 9 μg psPAX2 packaging plasmids, 6 μg VSV-G envelope plasmids and 2.5 M CaCl_2_ solution were mixed to a 50 mL Falcon tube and equal to 1.25 mL with Nuclease free water (Invitrogen). The DNA mixture was added dropwise into another 50 mL Falcon tube with fresh 1.25 mL 2X HBS (HEPES-buffered Saline) while vortex for efficient mixing. The reagent was then incubated at room temperature for 30 min. 2.5 mL of the mixture was dropped to the dish to ensure that the DNA mixture was evenly distributed on the dish's surface.

The transfection medium was removed from each dish, and 15 mL fresh cell line culture media with 10 mM Sodium Butyrate solution (Sigma-Aldrich) was added instead. After incubation at 37 °C, 5% CO_2_ for 8 h, media was harvested and kept at 4°C until the virus was concentrated. Another 20 mL fresh cell line culture media was replaced in the dish with the incubation at 37 °C, 5% CO_2_ overnight.

Media was collected from the dish, and all media was filtered by a 500 mL Bottle Top Vacuum Filter with 0.45 μm pore size. The filtered media was centrifuged at 6,000 g, 4 °C for 16-18 h in the J-25 Avanti centrifuge (Beckman), the F10BA-6x500y rotor (Beckman) with Beckman 500mL tubes (Beckman). After centrifugation, the supernatant was discarded carefully to not lose the pellet. 1-2 mL sterile cold DPBS (Dulbecco’s Phosphate Buffered Saline, Sigma-Aldrich) was gently added to wash the pellet. After washing, the DPBS washing pellet was replaced with the appropriate volume of fresh cold DPBS to dissolve the virus according to the size of the pellet. Once the pellet was suspended into DPBS, the solution with the virus was aliquoted and kept at -80 °C for storage before transduction.

Lentiviral transduction into primary cells and cell lines

For transduction into primary MECs, lentiviral particles were added into cells plated in 24-well suspension plates (Greiner Bio-One) with 0.1% polybrene (Sigma-Aldrich) and incubated at 37 °C, 5% CO_2_ for two days. After then, cells were pelleted by centrifuge and suspended in Matrigel for organoid culture.

Cell lines were transduced with lentiviral particles once cells cultured in the 35 mm dishes were grown to 80% confluency. Then 0.1% polybrene was added into the culture medium with lentiviral particles to increase the transduction efficiency. After 16 h, the medium was changed into fresh media before downstream experiments.

Organoid culture

Primary human MEC cells were suspended with 50 μL Matrigel and were seeded on 24-well suspension plates as domes (around 100,000 cells in Matrigel). Domes were then incubated at 37°C, 5% CO_2_ for 1 h to polymerize. 500 μL of organoid culture media were added into each well. Media was changed every three days, and Matrigel maintained for 6 days. After 6 days, one dome of organoids was passaged and spited into three domes seeded into new wells. For passaging, Matrigel was dissociated by 200 μL cell recovery medium, and the well plates were transferred and incubated on ice for 20 min. After Matrigel was degraded, organoids were centrifuged and pelleted for 5 min at 300 g at room temperature. Organoids were re-suspended by Matrigel, and domes were seeded again.

Circadian rhythms recording

LumiCycle (ACTIMETRCIS) recorded circadian rhythms in cells, which detects the bioluminescence of *Bmal1-*Luc in cells. Cells were seeded on 35 mm dishes and synchronized by 100 nM Dexamethasone (Sigma-Aldrich) for 1 h. After then, the culture medium changed to recording media. Dishes were then placed into the LumiCycle machine to trach circadian clocks.

To administer drugs, dishes were removed from LumiCycle. Drugs were directly put into recording medium. Dishes were then put back into LumiCycle to continue recording bioluminescence.

Time-series experiments

Cells were seeded on dishes and incubated till the confluence is reached to 70-80%. Then, cells were synchronized using 100 nM Dexamethasone for 1 h. After 24 h, cells were harvested every 3 h for 48h to extract protein and total RNA.

Hanging drop and cell invasion assay

Cells were trypsinized and resuspended at 400,000 cells/mL in hanging culture media with 0.24% methylcellulose (Sigma-Aldrich). 20 µL drops of cell suspension were seeded on the inside lid of the 10 cm culture dish. The lid was put back to the top of the dish so the drops were upside down. Spheroids were formed by gravity after incubation for 24 h at 37°C, 5% CO_2_. 10 µL hanging drops with spheroids were then transferred into a 48-well plate coated with 500 µL 1.8 mg/mL Collagen I mix. Notably, before seeding spheroids, Collagen I mix was placed at room temperature to settle down for 5 min. After seeding spheroid into Collagen I mix, the plates were placed into incubator at 37°C, 5% CO_2,_ to set for 2 h. Culture mediums with or without drug administration were added next. Cell invasions were captured daily under a snapshot microscope with Brightfield.

Generation of MCF-7/*ERα* KO cells by CRISPR-Cas9 editing

ERα was knocked out in MCF-7 cells by CRISPR-Cas9 system. Cas9 protein (Integrated DNA Technologies) and predesigned sgRNA (5’-TAGAGCGTTTGAYCATGAGC-3’) (Integrated DNA Technologies) were packaged by lipofectamine CRISPRiMax (Invitrogen) and transfected into MCF-7 cells with incubation at 37 °C for 2 days. After then, cells were trypsinized and seeded into 96-well plates for developing single colonies. When cells grew confluent, DNA was isolated from cells using ISOLATE II DNA genome kit (Meridian). CRISPR-edited DNA fragments were then amplified by PCR. PCR products were purified by QIAquick PCR Purification Kit and sequenced to select single colonies with *ERα* KO. The protein and mRNA expression levels were further examined to confirm *ERα* deletion in MCF-7 cells.

RNA isolation from human breast tissues

Human mammary gland tissues were placed into 1.5 mL Eppendorf tubes with RNA*later*^TM^ Stabilization Solution after surgery. Before isolating RNA, tissues were frozen at -80°C. Tissues were transferred to Lysing Matrix A with 1 mL cold QIAzol Lysis Regent supplemented from Rneasy Lipid Tissue Mini Kit and were disrupted and homogenized by TissueRuptor at 6 m/s, 30 s/cycle for 2 cycles. The homogenate was incubated at room temperature for 5 min and then centrifuged at 12,000 g for 5 min to remove the adipose layer on the top. Later, 200 μL chloroform was added along with shaking for 15 s, incubated again at room temperature for 3 min, and centrifuged at 12,000 g for 15 min at 4°C. The upper phase was then carefully transferred to a new 1.5 mL Eppendorf tube to avoid the middle white layer with DNA. Extra 200 μL fresh QIAzol was added, followed by shaking for 15 seconds, and incubated at room temperature for 1 min. Later, 200 μL chloroform was added to the mixture with incubation at room temperature for 1 min. Tubes were transferred to a centrifuge at 12,000 g, 4°C for 15 min. The upper phase was carefully taken as above and mixed with 185 μL participated sodium solution and 700 μL isopropanol after incubating at room temperature for 15 min. Then, the solutions were transferred into the Rneasy Mini spin column in a 2 mL collection tube, and RNA was extracted following QIAGEN RNeasy Mini Kit. The concentration of RNA was examined by Nanodrop 2000 software. For the RNA-seq experiment, the RNA quality was represented according to the RIN number checked using Tapestation 2200 (Agilent), and the concentration was tested using Qubit software (Life technology).

RNA to cDNA reverse transcription

Nanodrop determined the concentration of extracted RNA. 20 μL of each sample at 50 ng/μL concentration was added to the RT reaction tubes including RNA, water, RT Buffer Mix and RT Enzyme Mix in High-Capacity RNA to cDNA kit (Applied Biosystems). The RT reaction mix was placed into a Mastercycler at 37°C for 1 h and the enzyme was denatured at 95°C for 5 min. The cDNA was stored at -20°C and used within one month.

qPCR

10 ng cDNA was added into reaction wells consisting of 0.5 μL Taqman primers (Life technology) (see primer table), 4.3 μL Nuclease-free water, 5 μL Takyon ROX probe Mastermix DTTP blue (Eurogentec). Then plates were placed into a StepOne Plus RT-PCR system and cDNA was amplified by the protocol form StepOne v2.1 software (Applied Biosystems).

Histology of human breast tissues

Human tissues were fixed with 4% PFA (Paraformaldehyde, Sigma-Aldrich) for one day at 4°C, and then stored in 70% Ethanol at 4°C. Later, all samples were processed by Leica ASP300 Tissue processer overnight under a standard protocol. After processing, tissues were embedded in paraffin wax utilizing Leica EG1150H Embedding station.

Tissues were dissected into 5 μm thick pieces by S35 Microtome Blade on a Leica RM2255 Microtome. The sections were attached to Superfrost Plus slides. Later, slides were placed on a hot plate at 42°C and were transferred to a 42°C oven overnight to promote the adhesion to slides.

H&E staining

Tissue sections were dewaxed and rehydrated by dipping into Xylene for 3 min, 100% Ethanol for 2 min and 90% Ethanol for 2 min successively. Slides were stained using Hematoxylin for 5 min, washed by running tap water for 1 min and the residual Hematoxylin removed by dipping into 1% Acid Alcohol for 10 s. Following washing, Eosin-Y was used to counterstain tissue sections for 3 min before washing. Slides were dehydrated by a series of different concentrations of Ethanol (70%, 90% and 100%) and Xylene for 2 min of each step. Finally, the slides were mounted by DPX mountant and left in a fume hood overnight to dry.

Immunohistochemistry

Slides with tissue sections were dewaxed and rehydrated by sinking into xylene1 for 3 min, xylene 2 for 3 min, 100% Ethanol for 2 min, 90% Ethanol for 2 min and 70% ethanol for 2 min successively. Slides were then dipped into boiled Citrate Buffer 1X for 10 min to expose the antigen sites, left at room temperature for 20 min, and washed three times with 5 min intervals using PBS (Phosphate Buffered Saline). Later, sections were rinsed by PBS with 3% H_2_O_2_ (Hydroperoxide) at room temperature for 15 min to quench the activities of endogenous peroxidase in sections. After washing, the tissues were blocked by IHC blocking buffer at room temperature. Next, the primary antibody was added and incubated at 4 °C overnight.

The primary antibody was washed out using PBS for three times. The secondary antibody was then applied on tissue sections for 2 h. After washing steps, 1% streptavidin-horseradish peroxidase was added for 1 h at room temperature. After washing off unbound conjugate, DAB (Diaminobenzidine) solution was added, and then slides were counterstained by Hematoxylin solution for 30 sec. After successive dehydration following 70% ethanol for 3 min, 90% ethanol for 3 min, 100% ethanol for 3 min, xylene 1 for 5 min and xylene 2 for 5 min, slides were then mounted using DPX mountant. Lastly, slides were left in the fume hood overnight to dry. Images were captured by Slide Scanner, processed using CaseViewer and analyzed by ImageJ.

Immunocytochemistry

Cells were fixed using 4% PFA at room temperature for 30 min. Next, cells were permeabilized with 0.5% Triton X-100 in DPBS for 10 min. Cells were washed once using PBS for 10 min before incubation with blocking buffer that contains 10% Donkey serum for 1 h. After washing, primary antibodies were diluted in fresh blocking buffer and incubated at room temperature overnight.

In the following day, primary antibodies were washed out three time for 10 min each using DPBS. Secondary antibodies were then added and incubated at room temperature for 2 h in the dark. After washing, cells were stained for 0.001% Hoechst 33342 diluted into DPBS at room temperature for 15 min. Finally, the DPBS with Hoechst 33342 was replaced to fluorescent mounting media.

Western blotting

Cells were washed using cold DPBS for three times. RIPA lysis buffer with 1X protease inhibitor cocktail were added into dishes and left on ice for 10 min. Cells were then scraped and transferred into 1.5 mL Eppendorf tubes. Lysates were then centrifuged at 15,000 g, 4 °C for 20 min. Later, supernatants were transferred carefully into new 1.5 mL Eppendorf tubes for storage at -80 °C.

20 μg protein samples was diluted in 15 μL of water with 5 μL of 4X SDS sample loading buffer. The mixture was then centrifuged at 12,000 g for 1 min and denatured at 95 °C for 5 min. 15 μL of each sample was loaded into Mini-PROTEIN TGX gels and run at constant 110 V for 1 h in cold running buffer. Gels were transferred to nitrocellulose membrane using Trans-Blot Turbo Transfer System (Bio-Rad, CA, USA). After transfer, membranes were washed and protein was blocked in casein blocking buffer (Sigma-Aldrich) for 1 h at room temperature. Later, membrane was incubated with primary antibody diluted into blocking buffer overnight at 4 °C.

The membrane was washed three times in TBS-T (Tris Buffered Saline-Tween20) for 5 min each. Later, secondary antibody diluted into DPBS was added on the membrane with the incubation at room temperature for 1h. After repeating the washing steps as above, membranes were exposed using ECL (Enhanced Chemiluminescence) reagent.

Datasets of the hybrid design

The raw RSEM (RNA-Seq by Expectation Maximization) counts of all samples of breast invasive carcinoma cohort downloaded from FireBrowse (<http://firebrowse.org/>) were transformed to TPM values. The refGenome package (version 1.7.7) was used to parse the human gtf file (GRCh38). For a gene with multiple transcript splice-isoforms, we selected the longest transcript as the representative gene length. Raw reads of sequenced breast tumor and matched non-tumor adjacent samples from UK were aligned to human genome (GRCh38) using STAR (version 2.5)^3^. The expression quantification step was performed with RSEM (version 1.3.0)^4^. The raw RSEM counts were transformed to TPM values with similar steps as mentioned above. The rhythmic signals in transcriptome data can be disrupted by low sequencing depth. Thus, we only selected 26 non-tumor samples with sequencing depth above 20 million (> 20M) reads into the ordering step. For CYCLOPS ordering of non-tumor samples, we combined 26 time-stamped samples with those population samples from TCGA cohort and GTEx database without sampling time information. For further analysis of tumor samples with CYCLOPS, we only selected those samples collected from the centers that contribute to the non-tumor sample cohort.

Evaluate the clock function in sub-groups of population samples

Non-tumor and matched TCGA-BRCA tumor samples were selected based on the sample barcodes. Subtypes (Luminal A, Luminal B and Basal) of TCGA-BRCA tumor samples were extracted from previously published work^5^. R packages of RTCGA.clinical (20151101.20.0) and RTCGA (1.20.0) were used to 1) parse the clinical records of patients of breast invasive carcinoma cohort, 2) select ER+ and ER- tumor samples. The sequencing depth (> 20M reads) was used to select time-stamped tumor and non-tumor samples collected in UK. The correlation expression matrix of extracted sub-datasets of non-tumor or tumor samples were drawn with shiny app (https://github.com/gangwug/CCMapp).

**CYCLOPS 2.0 pre-processing**

Pre-processing methods were performed as described by Anafi et al. 2017^6^. Data from non-cancerous and Luminal A samples was only kept from batches with five or more non-cancerous samples. Probes were restricted to those in the “seed gene” list and the top 10,000 most highly expressed. Of these, the list was further limited to those with a coefficient of variation between 0.14 and 0.9. The upper bound of 0.9 was increased from 0.7, given the potential of batch and processing effects contributing to variance in aggregate data. For these probes, extreme expression values were capped at the top/bottom 2.5th percentile. The expression $X_{i,j}$ of each included probe $i$ in sample $j$ is scaled to give $S_{i,j}$:

$$S_{i,j}=\frac{X_{i,j}-M_{i}}{M_{i}},$$

where $M_{i}$ is the mean expression of probe $i$ across samples:

$$M_{i}=\left( \frac{1}{N} \right)\Sigma_{j}X_{i,j}.$$

The $S_{i,j}$ for non-cancerous data were expressed in eigengene coordinates $E_{i,j}$ following the methods of Alter et al. When non-cancerous and Luminal A data were combined, the $S_{i,j}$ data were expressed in the non-cancerous samples’ eigengene space. The number of eigengenes $N_{E}$ (singular values) retained was set to capture 85% of the seed data’s variance, as described by Anafi et al. 2017^6^.

**Covariate Processing**

Sample batch and tumor status were extracted from the expression files. Sample batch and tumor status were encoded into reduced one-hot flags. Reduced one-hot encoding resulted in an M-by-N matrix, where M is the sum of the number of groups in a covariate decreased by 1 for all covariates, and N is the number of samples in the dataset. This can be represented mathematically by $M=\sum_{c=1}^{C} {(m}_{c}-1)$, where C is the total number of covariates included, and m is the number of groups for covariate c. The non-cancerous data had seven batches, which resulted in a (7-1=) 6–by–j encoding of the covariate. The combined non-cancerous and Luminal A data included seven batches and two tumor types (non-cancerous vs Luminal A), which resulted in a (7-1+2-1) 7-by-N encoding of the covariates. To illustrate, an example is provided:

Two tumor types would conventionally be encoded as [1 0] (non-cancerous) and [0 1] (Luminal A) in one-hot representation. To simplify, the first index of the one-hot encoding is dropped, therefore encoding non-cancerous samples as [0] and Luminal A samples as [1]. In our analysis, this results in a 7-binary digit reduced one-hot, where the first 6 digits encode batch and the final digit encoded tumor status. A non-cancerous sample and Luminal A sample in batch one of seven are represented as [0 0 0 0 0 0 0] and [0 0 0 0 0 0 1], respectively. A non-cancerous sample and Luminal A sample in batch two of seven are represented as [1 0 0 0 0 0 0] and [1 0 0 0 0 0 1], respectively.

CYCLOPS 2.0 was optimized to fit the eigengene expression, including each sample's reduced one-hot covariate encoding.

**CYCLOPS 2.0 Model**

The CYCLOPS 2.0 Model comprises the *core structure* and the *covariate correction wrapper*. The *core structure* is the original CYCLOPS structure described in Anafi et al. 2017^6^ (Fig. S2A). The *covariate correction wrapper* has encoding and decoding layers, which linearly transform the sample eigengene expression on the encoding side. The decoding layer reverses this transformation. The encoding and decoding depend on the covariate group membership of a given sample. This can include one or more covariates such as batch, sex, or tumor status. The linear transformation consists of multiplicative (scaling) and additive terms. Specific scale and offset factors are fit for each covariate group (column) and eigengene (row). Prior to optimization, each eigengene's standard deviation and mean are calculated for each covariate group. The initial biases in the covariate encoding layer are chosen to be the average difference in eigengene expression between covariate groups. The initial scale factors in the covariate encoding layer are chosen to be the ratio of standard deviations in eigengene expression between covariate groups. The covariate wrapper's scale matrix and offset matrix have dimensions $N_{E}$-by-*M*, where $N_{E}$ is the number of eigengenes and M is the number of groups in a covariate, decreased by 1, summed across all covariates (in the previous example M=7).

We will use matrix notation to describe the CYCLOPS 2.0 model for simplicity. The first layer of the model is the encoding layer of the *covariate correction wrapper*, which performs a linear transformation depending on a covariate group membership.

$$covariate encoded data=E\circ\left( 1+S\cdot C \right)+B\cdot C+D$$

“$\circ$” is the element-wise matrix product. “$\cdot$” is the matrix dot product. The input, *E*, is an $N_{E}$-by-*one* column vector of the eigengene expression of sample *j*. *C* is an *M*-by-*one* column vector of the reduced one-hot covariate encoding of sample *j*. *S* is an $N_{E}$-by-*M* matrix for covariate group scale terms. *B* is an $N_{E}$-by-*M* matrix for covariate group offset terms. *D* is an $N_{E}$-by-*one* column vector of offsets added to all samples. The resulting *covariate encoded data* has the exact dimensions as the input, *E*.

*The covariate encoded data* is the input to the original CYCLOPS architecture, which consists of three layers. The first layer of the original architecture (and the second in the CYCLOPS 2.0 architecture) is a fully connected linear layer, which transforms the eigengene expression of a sample from $N_{E}$-dimensional space to two-dimensional space. The next layer of the original architecture, the third layer of the CYCLOPS 2.0 architecture, is the circular bottleneck layer, which performs a circular normalization. The final layer of the original architecture, and the fourth layer in the CYCLOPS 2.0 architecture, is another fully connected linear layer, which transforms the circular normalization from two-dimensional space back to $N_{E}$-dimensional space.

$$circular input=w_{in}\cdot covariate encoded data+b_{in}$$

$w_{in}$ is a *two*-by-$N_{E}$ matrix. $b_{in}$ is a *two*-by-*one* column vector. The *circular input* has two-by-one dimensions and is the input to the circular bottleneck function.

$$circular bottleneck output=\frac{circular input}{\sqrt{\sum{circular output^{\circ}}^{2}}}$$

“${^{\circ}}^{2}$” is the Hadamard, or elementwise, power. The denominator in the expression is the square root of the sum of squares of the *circular input*, represented by a single number. The division is performed elementwise. The *circular output* has the exact dimensions as the *circular input*. The sum of squares of the *circular bottleneck output* will always equal one for any given sample. A fully connected layer transforms the *circular output* from two-dimensional space back to $N_{E}$-dimensional space.

$$circular decoding=w_{out}\cdot circular output+b_{out}$$

$w_{out}$ is an $N_{E}$-by-*two* matrix: the inverse dimensions of $w_{in}$. $b_{out}$ is an $N_{E}$-by-*one* column vector.

The final layer of the model is the decoding layer of the *covariate correction wrapper*. This takes the *circular decoding* as the input and performs the inverse transformation of the *covariate correction wrapper*’s encoding layer.

$$covariate decoded data=\frac{circular decoding-(B\cdot C+D)}{1+S\cdot C}$$

The *covariate correction wrapper* has only one S and B matrix and one D column vector. The encoding and decoding layers of the *covariate correction wrapper* share these terms. This is intended to minimize the number of parameters and enforce the idea that these layers should be reflecting the same covariate-determined corrections. Notably, given the reduced one-hot encoding, there will always be a “base” group, where the covariate encoding is all zeros, and the expression data is left unchanged by the covariate encoding layer. All other groups are scaled relative to this base group. More formally, given the encoding used in this work $S\cdot C=0$ for non-cancerous samples in the first batch. Therefore, 1 is added to all $S\cdot C$, which makes the covariate scale term for non-cancerous samples in the first batch equal 1. Similarly, for non-cancerous samples in the first batch $B\cdot C=0$. Thus, non-cancerous samples in the first batch pass through the encoding and decoding layers of the *covariate correction wrapper* un-transformed.

In short, the algorithm performs the following transformations:

$$covariate encoded data=original Data\circ\left( 1+S\cdot C \right)+B\cdot C+D$$

$$circular bottleneck input=w_{in}\cdot covariate encoded data+b_{in}$$

$$circular bottleneck output=\frac{circular bottleneck input}{\sqrt{\sum{circular bottleneck input^{\circ}}^{2}}}$$

$$circular decoded data=w_{out}\cdot circular bottleneck output+b_{out}$$

$$covariate decoded model=\frac{circular decoded data-(B\cdot C+D)}{1+S\cdot C}$$

The colors show how the outputs of the previous function are provided to the following function. The goal is to minimize $\left( original Data-covariate decoded model \right)^{2}$, as will be explained.

**CYCLOPS 2.0 Ordering Algorithm**

The CYCLOPS 2.0 Model had to be optimized to obtain meaningful sample phase predictions. CYCLOPS 2.0, just like its forerunner, is an autoencoder algorithm. Its target is to recreate the input in the output. The input is the eigengene expression, and the output is the covariate decoded model. Thus, the loss function is defined as

$$\left( input-output \right)^{2}=\left( original Data-covariate decoded model \right)^{2},$$

known as the reconstruction error.

CYCLOPS 2.0 is implemented in julia 1.6, which offers a machine learning and optimization package, Flux.jl Flux.jl v0.13.5^7^. The “train!” function in Flux.jl was used to optimize the parameters of the CYCLOPS 2.0 model. The model was trained in two stages. In stage one, the ADAM optimizer^7^ was applied with a learning rate of 0.001 and a $\beta$ of (0.9, 0.999) for 1500 iterations. In stage two, the ADAM optimizer was applied with an initial learning rate of 0.001 and a $\beta$ of (0.9, 0.999), but if the reconstruction error increased from one iteration to the next, the optimization for that iteration was undone, and the learning rate was scaled by 0.5. If the reconstruction error was decreased from one iteration to the next, the learning rate was scaled by 1.1. This optimization was repeated for 2050 iterations or until the learning rate was reduced 1000-fold (equivalent to 10 consecutive learning rate decreases from the initial point), whichever came first. This is the Flux.jl implementation of the bold-drive optimization implemented in Anafi et. al 2017^6^.

Optionally, CYCLOPS 2.0 can include a secondary loss function, which uses sample collection time, if available for a small number of samples. This can help anchor the CYCLOPS ordering to an external time. When sample collection times are provided, the sum of the cosine distance between the predicted sample phase and the sample collection time is minimized in parallel to the reconstruction error. The collection time error is defined as

$$time balance\times\left( 1-\cos\left( CYCLOPS predicted phase-collection radian \right) \right)$$

A sequential search and 10-fold cross-validation determined the value of the *time balance* for this dataset. However, this term may vary for other datasets and the number of samples for which collection times are provided.

Parameter optimization was conducted for 80 random initial conditions and we selected the model that minimized the reconstruction error for a dataset. Parallelization was achieved using the package Distributed.jl, which includes the function “pmap.” Gradients were not explicitly defined but estimated by the “train!” function in Flux.jl.

Non-cancerous samples were first ordered independently. Luminal A samples were ordered in conjunction with the non-cancerous samples. The non-cancerous and Luminal A samples were projected in the non-cancerous samples’ eigengene space to emphasize the circadian variation in Luminal A samples, as was done in Anafi et al. 2017^6^.

**Obtaining CYCLOPS Sample Phase Predictions and Sample Magnitudes**

Sample phases are extracted as described by Anafi et al. 2017^6^. We calculate the angle between the positive x-axis and the coordinates represented by the *circular output* to obtain sample phase predictions. The *circular output* has two elements: an ‘x’ and a ‘y’ coordinate. The two elements in the *circular output* can be assigned ‘x’ or ‘y’ arbitrarily so long as this is chosen consistently throughout all analyses. To transform the *circular output* to an angle, we use the *atan* function in julia, $\mathrm{atan}\left( y,x \right)$, essentially the same as the $\tan^{-1} \left( y/x \right)$. However, the *atan* function assigns the correct angle in all coordinate quadrants. Of some note, the *circular input* could be used instead of the *circular output*, as the ratio of ‘x’ and ‘y’ coordinates is the same within the two sets of coordinates; using the *circular output* is simply a convention.

The CYCLOPS sample magnitude (CMag) is the distance of the *circular input* from the inferred origin (0, 0) (Fig. 5A); this is the square root of the sum of squares of the *circular input*. Thus, the magnitude of a sample is given by:

$$magnitude=\sqrt{\sum(circular input{^{\circ}}^{2})}=\sqrt{x^{2}+y^{2}}.$$

**Assessing Transcript Rhythmicity**

Transcript rhythmicity was assessed using modified cosinor regression as performed by Anafi et al. 2017^6^, where CYCLOPS 2.0 gave predicted sample phases. Cosinor regression is a statistical method of fitting a sinusoidal function to data^8^. These data are assumed to be the result of a periodic process, and this approach works best when data are provided for multiple cycles of a process. CYCLOPS assigns sample phases between $0$ and $2\pi$, covering only a single cycle. We modify the standard cosinor regression approach to avoid misidentifying a monotonic process as a cyclic process.

First, we fit a linear regression line to the expression of a single transcript:

$$g_{i,j}=l_{i}\left( \Theta_{j,i} \right)+c_{i}+\epsilon_{i}\ldots\left[ M1 \right],$$

where

$$\Theta_{j,i}=\left( \theta_{j}-\phi_{i} \right)\%2\pi\ldots\left[ M2 \right],$$

and $\theta_{j}$ is the raw CYCLOPS predicted phase for sample j.

The formula M1 states that the expression of a transcript i in sample j is modeled by $l_{i}$ times the phase of that sample $\Theta_{j,i}$. The formula M2 states that for each transcript i, all sample phases $\Theta_{j,i}$ are shifted by a common term $\phi_{i}$ and the modulus (%) of the sample phases divided by $2\pi$ is calculated. A brute force search optimizes the term $\phi_{i}$. The ideal shift for each transcript $\phi_{i}$ minimizes the difference between transcript expression and the predicted expression of transcript i in sample j, as compared to all other tested shifts. The formula M1 becomes the null hypothesis for the cosinor regression.

Once the ideal shift $\phi_{i}$ for the linear regression line has been established, we next fit a line with both monotonic and cyclic parameters:

$$g_{i,j}=l_{i}\left( \Theta_{j,i} \right)+\alpha_{i}*\cos\left( \Theta_{j,i} \right)+\beta_{i}*\sin\left( \Theta_{j,i} \right)+c_{i}+\epsilon_{i}\ldots[M3]$$

To get the regression f-statistic, we first calculate the residual sum of squares given by M1 and M3 ($SSE_{M1}$ and $SSE_{M3}$, respectively). Next, we can calculate:

$$F_{i}=\frac{\left( SSE_{M1}-SSE_{M3} \right)/2}{{SSE_{M3}}/\left( N-t \right)},$$

where N is the number of samples in the expression data, and t is the number of parameters in the model M3. When only one batch of expression data is present, $t=3$. For each additional batch, t increases by 1.

We use the ‘FDist’ function from the ‘Distributions.jl’ package in julia 1.6^9^ to generate an F-distribution. We use the F-distribution and the ‘cdf’ function to calculate the cumulative probability at $F_{i}$. We can calculate the p-statistic of each transcript by subtracting the transcripts' calculated cumulative probability from one. To avoid type 1 error, we calculate the ‘Benjamini-Hochberg’ multiple test correction q-value from all transcript p-values.

Lastly, we fit a cosinor model without monotonic terms:

$$g_{i,j}=\alpha_{i}*\sin\left( \theta_{j,i} \right)+\beta_{i}*\cos\left( \theta_{j,i} \right)+c_{i}+\epsilon_{i}\ldots\left[ M4 \right].$$

The predicted phase of peak expression of transcript, $i$, is given by:

$$\Phi_{i}=\mathrm{atan} \left( \alpha_{i},\beta_{i} \right)\%\left( 2\pi\right).$$

The predicted amplitude of the cosinor of transcript i is given by:

$$Amp_{i}=\sqrt{\alpha_{i}^{2}+\beta_{i}^{2}}.$$

The predicted average expression of transcript, $i$, is given by $c_{i}$. CYCLOPS 2.0 orders multiple batches of data and therefore includes batch as an indicator for $l_{i}, \alpha_{i}, \beta_{i},$ and $c_{i}$ in $M1, M3,$ and $M4$. The F-statistic is evaluated for all samples across all batches, not by batch.

**Aligning CYCLOPS Predicted Phases to Known Biology**

The sample phase predictions produced by CYCLOPS reflect a relative ordering. A circle has no intrinsic beginning or endpoint. Additional information is required to relate the ordering to an external reference e.g., circadian time 0, wall clock time 4 PM, or dim light melatonin onset (DLMO).

To align sample phases to known biology, we use the predicted sample phases to calculate the cosinor of best fit for 17 mouse atlas genes^10^. Using CYCLOPS-predicted sample phases, these genes' predicted peak expression times are aligned with their previously established peak circadian expression times averaged across mouse tissues. This is similar to what was done in Anafi et al. ^6^, where the average acrophase of the PAR bZip genes was set to pi, and Rubin et al.^11^ where the acrophase of ARNTL was set to 0. Here, rather than using a single gene, the alignment was set to best match the entire core clock repertoire seen in mice.

A brute force search finds the optimal alignment between predicted and established gene peak expression times. The direction and shift that minimizes the mean cosine distance between predicted and established peak expression times is applied to all predicted sample phases. This gives the ‘mouse atlas circadian time’-aligned predicted sample phases.

To determine which of the 17 genes are used for the alignment, we first calculate their cosinor regression p-statistic (explained in ‘Assessing Transcript Rhythmicity’). Of the 17 genes, only those with a *p*-statistic < 0.05 are considered for the alignment. We cannot define peak expression times in our data ordering for arhythmic genes.

**Gene Set Enrichment Analysis (GSEA) for Hallmark Gene Sets Enriched for Cycling Transcripts in Non-cancerous Tissue**

Enrichment of MSigDB Hallmark Gene Sets by cycling transcripts in non-cancerous tissue was assessed in GSEAPreranked v4.2.3^12^. The natural log of the cosinor regression f-statistic was used to rank transcripts. The enrichment statistic was set to “classic.” All other “Basic” and “Advanced” fields were set to default values.

**Phase Set Enrichment Analysis (PSEA) for Hallmark Gene Sets Phase Coordinated among Cycling Transcripts in Non-cancerous and Luminal A Tissue**

PSEA^13^ was separately applied to cycling transcripts in non-cancerous and Luminal A tissue to test if they were phase coordinated in MSigDB Hallmark gene sets. In each case, only transcripts with a cosinor regression BHq<0.05 and a relative amplitude (amplitude/MESOR) > 0.33 were included in the lists. The domain maximum was set to $2\pi$ since acrophases were provided in radians. All other fields were left as defaults. Only gene sets with a BHq<0.05 were shown in the graphic.

**Overrepresentation Analysis using EnrichR of Significantly Cycling Transcripts in Non-cancerous Tissue in Hallmark Gene Sets**

Transcripts cycling with a BHq<0.05 and relative amplitude>0.33 were included in this list. EnrichR v3.1^14^ in R was used to test for overrepresentation of MSigDB Hallmark gene sets by these cycling transcripts.

**GSEA for Amplitude Change between Luminal A and Non-cancerous Samples**

First, the list of transcripts was restricted to those cycling in non-cancerous or Luminal A tissue with a BHq<0.05. Next, a cosinor regression model was fit to transcript expression that included tissue type-specific amplitude terms, and batch offset terms. This cosinor model can be expressed as:

$$g=A*sin \left( \theta\right)+B*cos \left( \theta\right)+A_{C}*I*\sin\left( \theta\right)+B_{C}*I*\cos\left( \theta\right)+C+C_{C}*I+batch,$$

Where the indicator I is set to 0 for non-cancerous samples and 1 for Luminal A samples. Therefore, for non-cancerous samples, the model simplifies to:

$$g=A*sin \left( \theta\right)+B*cos \left( \theta\right)+C+batch.$$

The transcript amplitude of the non-cancerous tissue is $Amp_{nc}=\sqrt{A^{2}+B^{2}}$, and the transcript amplitude for the Luminal A tissue is $Amp_{LumA}=\sqrt{\left( A+A_{C} \right)^{2}+\left( B+B_{C} \right)^{2}}$. The natural log of the amplitude ratios, $\ln\left( \frac{Amp_{LumA}}{Amp_{nc}} \right)$, were calculated for these genes and used to rank transcripts. This list was supplied to GSEAPreranked. The enrichment statistic was set to “classic.” All other “Basic” and “Advanced” fields were left at default values. Positive enrichment means increased amplitude in Luminal A tissue, whereas negative enrichment means increased amplitude in non-cancerous tissue.

**EnrichR for Differential Cycling Amplitude between Luminal A and Non-cancerous Samples**

Again, transcripts were restricted to significantly cycling in non-cancerous or Luminal A tissue with a BHq<0.05. The significance of differential cycling was now assessed using nested cosinor regression. The base model (null hypothesis) for the differential cycling regression is:

$$g=A*sin \left( \theta\right)+B*cos \left( \theta\right)+C+C_{C}*I+batch,$$

and the expanded model (alternative hypothesis) is:

$$g=A*sin \left( \theta\right)+B*cos \left( \theta\right)+A_{C}*I*\sin\left( \theta\right)+B_{C}*I*\cos\left( \theta\right)+C+C_{C}*I+batch.$$

An F-test was used to assess the null hypothesis $A_{C}=B_{C}=0$. Transcripts showing differential cycling with a BHq<0.05 were identified. To test for increased amplitude in Luminal A tissue, we further restricted significantly differentially cycling transcripts to those with $\frac{Amp_{LumA}}{Amp_{nc}}>5$. To test for increased amplitude in non-cancerous tissue, we restricted significantly differentially cycling transcripts to those with $\frac{Amp_{LumA}}{Amp_{nc}}<0.1$. These lists were provided to EnrichR to test the overrepresentation of increased and decreased cycling in MSigDB Hallmark gene sets.

**CYCLOPS Magnitude–Transcript Amplitude Analysis in Luminal A Samples**

The CMags of all Luminal A samples were gathered and binned into three equally sized groups: low, medium, and high (Fig. 5B). Only samples in the *low* and *high* groups were retained for this analysis. Cosinor regression was performed with the model defined as:

$$g=A*\sin\left( \theta\right)+B*\cos\left( \theta\right)+A_{high}*I_{high}*\sin\left( \theta\right)+B_{high}*I_{high}*\cos\left( \theta\right)+C+batch.$$

The amplitude of the cosinor fit of the low-CMag samples is calculated by:

$$Gene Amp_{low}=\sqrt{A^{2}+B^{2}}.$$

The amplitude of the cosinor fit of the high-CMag samples is calculated by:

$$Gene Amp_{high}=\sqrt{\left( A+A_{high} \right)^{2}+\left( B+B_{high} \right)^{2}}.$$

The natural log ratio of the high-CMag amplitude over the low-CMag amplitude was calculated for all significantly cycling transcripts in Luminal A (Fig. 5C):

$$Mag Amp Ratio=\ln\left( \frac{Gene Amp_{high}}{Gene Amp_{low}} \right).$$

**CYCLOPS Magnitude-Prognosis Analysis in Luminal A Samples**

To test if patients had different likelihoods of bad outcomes by CMag group, we first fit a generalized linear model to patient death within five years of diagnosis as a function of age, metastasis at diagnosis, and CMag group. We then fit these data to a logistic regression model in R using the glm function and the binomial linker function. This full model is written as follows:

$$death_{<5 years}=age+mets_{diagnosis}+CMag_{group},$$

and is compared to the partial model:

$$death_{<5 years}=age+mets_{diagnosis}.$$

The significance of $CMag_{group}$ term is assessed by nested Chi-squared test using the ‘pchisq’ function in R v4.2.1, and setting ‘lower.tail=F’.

**GSEA for Differential Cycling Amplitude within Luminal A Samples, between High- and Low-CMag**

Transcripts were restricted to those significantly cycling in Luminal A with BHq<0.05. We computed the ratio of the amplitude of each transcript in the high magnitude group as compared to the amplitude of the same transcript in the low magnitude group. We ranked the transcripts by the natural log (ln) of these amplitude ratios. The ranked listed was analyzed in GSEA using the “preranked” function to test for enrichment of MSigDB Hallmark gene sets. The enrichment statistic was set to “classic.” All other “Basic” and “Advanced” fields were left as defaults.

**EnrichR for Differential Cycling Amplitude within Luminal A Samples, between High- and Low-CMag**

Transcripts were restricted to those significantly cycling in Luminal A with a BHq<0.05. These transcripts were limited to significantly differential amplitudes between High- and Low-CMag groups with a BHq<0.05. This list was provided to enrichR to test for the overrepresentation of significant cycling and significantly differential transcripts in MSigDB Hallmark gene sets. No transcripts were identified at BHq<0.05 that had a higher amplitude in the low CMag samples.

**CYCLOPS Benchmarking Using BA11 Dataset and Observed Differences between Two Lung Datasets**

We sought to evaluate our ability to reconstruct rhythms in the setting of realistic interindividual differences in rhythmic transcript expression and realistic differences in batch processing. We used the differences observed between the Groningen and Laval lung datasets^15^ to model the differences in cycling transcript expression that are expected to arise from batch processing. We used data describing transcript expression in the pre-frontal cortex (BA11) as a base model of rhythmic transcript expression^16^. These BA11 cortical data were obtained at a single center and spanned the entire circadian cycle. We aimed to generate semi-synthetic data to approximate the BA11 expression observed at a theoretical second center, including batch processing and sample collection time differences. We sequentially increased the complexity of our synthetic data model. We evaluated the performance of both the original algorithm and CYCLOPS 2.0 at each step (Fig. S2B-D).

Initially, to simply model the differences expected to arise as a result of batch processing, we calculated the mean and standard deviation of every transcript $i$ in both Groningen and Laval data, $M_{G,i}$ and $M_{L,i}$, and $\sigma_{G,i}$ and $\sigma_{L,i}$. Then, we calculated the ratios of Groningen to Laval parameters for each transcript, $M_{R,i}={M_{G,i}}/{M_{L,i}}$ and $\sigma_{R,i}={\sigma_{G,i}}/{\sigma_{L,i}}$. We observed a linear correlation between these parameter ratios:

$$\bar{M_{R,i}} \sim0.31\times\sigma_{R,i}+\epsilon_{M,i},$$

where ‘$\bar{M_{R,i}}$’ denotes the predicted value of $M_{R,i}$, and $\epsilon_{M,i}$ denotes the residuals about this line. Based on the observed parameter distributions, we modeled the residuals $\epsilon_{M,i}$ using a log-normal distribution:

$$\bar{\epsilon_{M,i}} \sim e^{N\left( \mu=-0.33,\sigma=0.24 \right)}.$$

Similarly, we modeled the standard deviation ratio using a log-normal distribution:

$$\bar{\sigma_{R,i}} \sim e^{N\left( \mu=0.5,\sigma=0.21 \right)}.$$

We then redefined the $\bar{M_{R,i}}$ model such that:

$$\bar{M_{R,i}} \sim\bar{\sigma_{R,i}}\times0.31+\bar{\epsilon_{M,i}}+\epsilon_{i}.$$

With these three formulations, we can generate random mean and standard deviation ratios that mimic those observed comparing the Groningen and Laval data sets. First, we scaled the spread of each transcript about its mean. We computed the mean of each transcript in the BA11 dataset across all samples. We subtracted the mean value for each transcript in every sample, giving us the differences between each observation and the mean value for each transcript. We then scaled these data, multiplying each row (transcript) by a random number drawn from $\bar{\sigma_{R,i}}$. Next, we adjusted the mean expression of each transcript. We drew random numbers from $\bar{M_{R,i}}$ to scale mean transcript expression. This gives us the desired scaled mean $\mu_{i}\times\bar{M_{R,i}}$. After combining the scaled sample mean value and scaled sample spread, we added random noise following a normal distribution with a mean of 0 and a standard deviation of 5% of the expression average of a transcript. These data reflect a model of BA11 expression data that might be observed at a theoretical second center. Finally, we combined the original BA11 expression data with the modeled second center to give a semi-synthetic dataset with observed mean and standard deviation ratios between batches, as observed between Groningen and Laval.

To make the mean and standard deviation ratios between batches more or less pronounced, we can exponentiate the log-normal distributions of $\bar{M_{R,i}}$, and $\bar{\sigma_{R,i}}$ by $n$. To make the two batches more like each other, $n<1$, and to make batches less like each other, $n>1$; we can draw mean and standard deviation ratios from ${\bar{M_{R,i}}}^{n}$ and ${\bar{\sigma_{R,i}}}^{n}$.

For our benchmarking, we tested $n\in\{0.0, 0.1, 0.5, 1.0, 2.0\}$. For each $n$, we generated five replicates of semi-synthetic datasets. Batch ids were added to the data for CYCLOPS 2.0. We initialized the ordering process from 80 different random starting positions and selected the original and CYCLOPS 2.0 models that minimized the reconstruction error for a dataset. We assessed their average sample angular error for each reconstruction (Fig. S2B).

**ComBat Normalization**

We also intended to compare CYCLOPS 2.0 performance to the performance of the original CYCLOPS on ComBat normalized data^17^. COMBAT is an established RNA-seq batch normalization method. We used ComBat in R v4.2.1 provided by the sva.R v3.44.0 package^18^.

To normalize batch data, all expression data must be greater than 0. As it was possible that one of the randomly generated expression values might be below 0, the minimum value of the entire dataset minus one was subtracted from all values:

$${expr}_{1+}={expr}_{i,j}-\min\left( expr \right)+1.$$

This made every value in the dataset greater than or equal to one. This positive expression data, ${expr}_{1+}$, were log-transformed before being provided to combat:

$${expr}_{ln}=\ln\left( {expr}_{1+} \right).$$

The combat-normalized data were unlogged and translocated in the opposite direction as previously:

$$expr_{norm}=e^{expr_{ComBat}}+\min\left( expr \right)-1.$$

**CYCLOPS Benchmarking Using BA11 Dataset and Observed Differences between Two Lung Datasets with Sample Time Bias**

In our first benchmark test, the synthetic and semi-synthetic data had the same distribution of sample collection times. Next we aimed to benchmark the performance of the original CYCLOPS and CYCLOPS 2.0 in the presence of in sample collection time biases. While one collection site was modeled to include samples from across the circadian period, the other was restricted to a defined temporal window. Notably, these differences are expected in our application as GTEx is an autopsy-based collection while TCGA data is collected during clinical care. We again also modeled transcript expression differences that arise from batch processing.

We first used CYCLOPS to separately order the Groningen and Laval datasets and then assessed the rhythmicity of each transcript, $i$, in both datasets. Rhythmicity was assessed as explained in ‘Assessing Transcript Rhythmicity.’ We categorized each transcript as either cycling in both datasets or not. To clarify, if a transcript was cycling in one dataset but not the other, it was categorized as not cycling.

For all non-cycling transcripts, we calculated the mean and variance of every transcript, $i$, in both Groningen and Laval data, $M_{G,i}$ and $M_{L,i}$, and $\sigma_{G,i}^{2}$ and $\sigma_{L,i}^{2}$. We calculated the ratio of means in Groningen to Laval for each transcript, $i$, $M_{R,i}={M_{G,i}}/{M_{L,i}}$, and modeled these ratios using a log-normal distribution:

$$\bar{M_{R,i}} \sim e^{N\left( \mu=0.16,\sigma=0.16 \right)}.$$

Instead of calculating the standard deviation ratios, we calculated the ratio of transcript standard deviations relative to transcript mean ratios between datasets:

$$\sigma_{R,i}^{2}=\frac{\left( {\sigma_{G,i}^{2}}/{\sigma_{L,i}^{2}} \right)}{M_{R,i}}.$$

We modeled the relative standard deviation ratios using a log-normal distribution:

$$\bar{\sigma_{R,i}^{2}} \sim e^{N\left( \mu=0.96,\sigma=0.53 \right)}.$$

For each cycling transcript, we calculated the MESOR ($B_{G,i}$ and $B_{L,i}$), amplitude ($A_{G,i}$ and $A_{L,i}$), and residual variance ($\delta_{G,i}^{2}$ and $\delta_{L,i}^{2}$), in each dataset. We start by calculating the MESOR ratio:

$$B_{R,i}={B_{G,i}}/{B_{L,i}},$$

and model the ratio using a log-normal distribution:

$$\bar{B_{R,i}} \sim e^{N\left( \mu=0.21,\sigma=0.43 \right)}.$$

We calculate the ratio of transcript amplitude relative to the transcript mean ratio:

$$A_{R,i}=\frac{\left( {A_{G,i}}/{A_{L,i}} \right)}{M_{R,i}},$$

and model the ratio using a log-normal distribution:

$$\bar{A_{R,i}} \sim e^{\left( \mu=0.34,\sigma=0.31 \right)}.$$

We calculate the ratio of transcript variance relative to the transcript amplitude ratio:

$$\delta_{R,i}^{2}=\frac{\left( {\delta_{G,i}^{2}}/{\delta_{L,i}^{2}} \right)}{A_{R,i}\times M_{R,i}},$$

and model the ratio using a log-normal distribution:

$$\bar{\delta_{R,i}^{2}} \sim e^{N\left( \mu=0.44,\sigma=0.49 \right)}.$$

To briefly recap, we have modeled the mean ratio, $\bar{M_{R,i}}$, and relative variance ratio, $\bar{\sigma_{R,i}^{2}}$, distributions for non-cycling transcripts, and the MESOR ratio, $\bar{B_{R,i}}$, relative amplitude ratio, $\bar{A_{R,i}}$, and relative residual variance, $\bar{\delta_{R,i}^{2}}$, distributions for cycling transcripts. These distributions reflect the scale and offset ratios between Groningen and Laval.

We aim to create synthetic BA11 data from a hypothetical clinical center to integrate with the existing autopsy based BA11 expression data. This enables us to validate the predicted phase orderings in the presence of both expression differences and sample phase collection biases between batches. Using the calculated ratios between Groningen and Laval, we can generate synthetic BA11 parameters from the real BA11 parameters. For each *non-cycling* transcript, $i$, in the BA11 data, we calculate the mean, $M_{i}$, and variance, $\sigma_{i}^{2}$. For each *cycling* transcript, $i$, we calculate the acrophase, $\Phi_{G,i}$, MESOR, $B_{i}$, amplitude, $A_{i}$, and residual variance, $\delta_{i}^{2}$. The synthetic mean of a non-cycling transcript, $i$, is given by:

$$M_{S,i}=\frac{M_{i}}{\bar{M_{R,i}}} ,$$

and the synthetic transcript variance of a non-cycling transcript, $i$, is given by:

$$\sigma_{S,i}^{2}=\frac{M_{i}}{\bar{M_{R,i}}\times\bar{\sigma_{R,i}^{2}}} .$$

The synthetic MESOR of a cycling transcript, $i$, is given by:

$$B_{S,i}=\frac{B_{i}}{\bar{B_{R,i}}} ,$$

the synthetic amplitude by:

$$A_{S,i}=\frac{A_{i}}{\bar{B_{R,i}}\times\bar{A_{R,i}}} ,$$

and the synthetic residual variance by:

$$\delta_{S,i}^{2}=\frac{\delta_{i}^{2}}{\bar{B_{R,i}}\times\bar{A_{R,i}}\times\bar{\delta_{R,i}^{2}}} .$$

To generate synthetic expression data for a non-cycling transcript $i$, in sample $j$, we draw random numbers from a normal distribution with a mean $M_{S,i}$ and variance $\sigma_{S,i}^{2}$:

$${non-cycling expression}_{i,j} \sim N\left( \mu=M_{S,i},\sigma=\sqrt{\sigma_{S,i}^{2}} \right).$$

To generate synthetic expression data for a cycling transcript, $i$, we first need to draw random sample collection times. For this purpose, we drew random sample collection times from five windowed uniform distributions; 24-, 12-, 10-, 8-, and 6-hour windows (Fig. S2E ’12-hour window’ – ‘uniform’). We drew 101 random sample collection times, $\theta_{j}$, from each windowed uniform distribution five times. Each dataset comprises 202 real BA11 samples and 101 synthetic BA11 samples. The synthetic expression data for a cycling transcript $i$, in sample $j$, is given by:

$${cycling expression}_{i,j} \sim A_{S,i}\times\cos\left( \theta_{j}-\Phi_{i} \right)+M_{S,i}+N\left( \mu=0,\sigma=\sqrt{\sigma_{S,i}^{2}} \right).$$

We combine the non-cycling and cycling expression data to give a model of BA11 expression data. Finally, we integrate the synthetic expression data with the real BA11 expression data. Batch identifiers were added to the data for CYCLOPS 2.0 training.

For the original model, synthetic expression data were normalized by batch identifier group using ComBat as outlined in ‘ComBat Normalization.’ Each synthetic dataset has a ComBat-normalized pair. CYCLOPS was used to order each of the 25 synthetic ComBat-normalized datasets. CYCLOPS 2.0 models was used to order each of the 25 synthetic datasets without normalization. In both cases we initialized the ordering process from 80 different random starting positions and selected the original and CYCLOPS 2.0 models that minimized the reconstruction error for a dataset. These synthetic data have known sample collection phases. We assessed the average sample angular error using both ordering approaches. (Fig. S2C).

**CYCLOPS Benchmarking Using Observed Differences between Two Lung Datasets with Sample Time Bias**

The number of cycling transcripts in brain tissue has been found to be lower than the number of cycling transcripts in liver, lung, and kidney^10,11^. The amplitude of individual cycling transcripts in the brain tissues also tended to be lower. In our experience ordering of brain tissues tends to be a more difficult challenge. Thus, we also sought to compare the performance of CYCLOP 2.0 and CYCLOPS with ComBat correction in a tissue with stronger cycling. We repeated the above steps but imitated the cycling parameters of the lung datasets^15^. The *non-cycling* mean, $M_{i}$, and variance, $\sigma_{i}^{2}$, and *cycling* acrophase, $\Phi_{i}$, MESOR, $B_{i}$, amplitude, $A_{i}$, and residual variance, $\delta_{i}^{2}$, where calculated from Groningen data. The sample collection times for the synthetic Groningen data were drawn from a random uniform distribution ranging from $0$ to $2\pi$. The cycling and non-cycling parameters for the synthetic Laval data were generated from the modeled ratio distributions above and the Groningen parameters, following the steps in the previous section. For the synthetic Laval dataset, sample collection times were drawn from the 5 windowed distributions (Fig. S2E ‘12-hour window’ – ‘uniform’) in replicates of 5.

Each dataset contained 400 samples, 300 synthetic Groningen samples, and 100 synthetic Laval samples. Samples were labeled by batch for CYCLOPS 2.0. Datasets were ComBat normalized by batch identifier, as outlined above. CYCLOPS was used to order each of the 25 synthetic ComBat normalized datasets. CYCLOPS 2.0 was used to order the each of the 25 synthetic datasets without normalization but including a batch identifier label. In both cases we initialized the ordering process from 80 different random starting positions and selected the original and CYCLOPS 2.0 models that minimized the reconstruction error. These synthetic data have known sample collection phases. We assessed the average sample angular error using both ordering approaches. (Fig. S2D).

**Quantification and statistical analysis**

All grouped data were presented as Mean ± SEM with standard nomenclature for significance values, **p<*0.05, ***p<*0.01, ****p<*0.001, *****p<*0.0001. Data values and variance were calculated as the average of all experimental repeats. All figures were made by GraphPad Prism 9 software and R, except for the bioluminescent recording experiment, which was created by LumiCycle Analysis software (ACTIMETRICS). The specific statistic tests used were specified in figure legends. The Fig. 1A was created with [BioRender.com](https://biorender.com/).
