## Supplemental Information for "Subtype-specific circadian clock dysregulation modulates breast cancer biology, invasiveness, and prognosis"

**Table. S1 Number of Batch Processing Sites, Samples, and Patient Average Ages for Non-Cancerous Female Breast Samples (Related to Fig. 2)**

| Data Source | GTEx | TCGA | UK |
| --- | --- | --- | --- |
| Number of Batch Processing Sites | 2 | 4 | 1 |
| Number of Samples | 167 | 106 | 26 |
| Average Age (mean) | 50.7 | 56.7 | 57.9 |

**Table. S2 Number of Batch Processing Sites, Samples, and Patient Average Ages for Luminal A Female Breast Samples (Related to Fig. 4)**

| Source | TCGA | UK |
| --- | --- | --- |
| Number of Batch Processing Sites | 4 | 1 |
| Number of Samples | 193 | 18 |
| Average Age (mean) | 58.8 | 59.6 |

**Supplementary 2,**

**File. S1 Cycling Parameters Describing Significantly Cycling Transcripts in Non-Cancerous Female Breast Tissue (Related to Fig. 2)**

This file includes the cycling parameters estimated for significantly cycling transcripts in non-cancerous samples. CYCLOPS 2.0 was used to order non-cancerous GTEx, TCGA, and UK (Manchester) female breast samples. Transcript rhythmicity was assessed by modified cosinor regression (see methods). Cycling transcripts were defined by a FDR (BH q-statistic) < 0.05, and a relative amplitude (amplitude over MESOR) > 0.33. For each cycling transcript we report the *MESOR*, *Amplitude*, *Amplitude over MESOR*, *Acrophase*, modified cosinor regression *p-statistic*, *q-statistic*, and *f-statistic*.

**Supplementary 3,**

**Video. S1 Representative LV200 bioluminescence recording in human tumor and normal organoids isolated from Luminal A patient (Related to Fig. 3C)**

**Supplementary 4,**

**Video. S2 Representative LV200 bioluminescence recording in human tumor and normal organoids isolated from TNBC patient (Related to Fig. 3D)**

**Supplementary 5,**

**File. S2 Cycling Parameters Describing Significantly Cycling Transcripts in Luminal A Female Breast Tissue (Related to Fig. 4)**

This file includes the cycling parameters estimated for significant cycling transcripts in Luminal A samples. CYCLOPS 2.0 was used to order non-cancerous and Luminal A samples from TCGA and the UK (Manchester). Transcript rhythmicity in the Luminal A samples was assessed by modified cosinor regression (see methods). Cycling transcripts were defined by an FDR (BH q-statistic) < 0.05, and a relative amplitude (amplitude over MESOR) > 0.33. For each cycling transcript we report the *MESOR*, *Amplitude*, *Amplitude over MESOR*, *Acrophase*, modified cosinor regression *p-statistic*, *q-statistic*, and *f-statistic*.

**Supplementary 6,**

**File. S3 Differential Cycling Analysis Comparing Luminal A Relative to Non-Cancerous Female Breast Tissue (Related to Fig. 4)**

Nested cosinor regression models (see methods) were used to assess the significance of differential cycling comparing non-cancerous and Luminal A samples. All samples that were independently identified as rhythmic in either Luminal A or non-cancerous samples (BH q-statistic < 0.05) are included in this analysis. For each transcript we report the significance of differential rhythmicity as assessed by nested cosinor regression p-statistic and q-statistic along with the ratio of transcript amplitude in Luminal A as compared to non-cancerous samples.

**Supplementary 7-9,**

**File S4-6. GSEA Results for Therapeutic-Related Gene Sets Positively Enriched for Cycling in Luminal A (Related to Fig. 4)**

Transcripts were ranked by the modified cosinor regression f-statistic assessing rhythmicity in Luminal A samples. Pre-ranked GSEA was used to identify therapeutic related genes sets enriched for cycling. Results are provided for gene sets available in DsigDB D1 FDA Approved Drugs^1^, Kinase Perturbations from GEO Down, and Kinase Perturbations from GEO Up.

**Supplementary 10,11,**

**File S7, S8 Enrichr Results for Therapeutic Related Gene Sets Overrepresented for Significant Cycling Transcripts in Luminal A (Related to Fig. 4)**

Enrichr was used to identify therapeutic related gene sets that showed overrepresentation of transcripts cycling in Luminal A samples. Transcripts with a BHq < 0.05 and relative amplitude (amplitude/MESOR) > 0.33 were included. Results are provided for gene sets included in Kinase Perturbations from GEO Down, and Kinase Perturbations from GEO Up.

**Supplementary 12,**

**Video. S3 Representative Incucyte live imaging for determining proliferation of *BMAL1-*KD MCF-7 cells**

1.	Yoo, M., Shin, J., Kim, J., Ryall, K.A., Lee, K., Lee, S., Jeon, M., Kang, J., and Tan, A.C. (2015). DSigDB: drug signatures database for gene set analysis. Bioinformatics *31*, 3069-3071. 10.1093/bioinformatics/btv313.
